# Multiplexed CRISPR/Cas9 Editing of Tumor Suppressor Genes Recapitulates Molecular and Morphological Features of High-Risk Endometrial Cancer

**DOI:** 10.1101/2025.11.28.691175

**Authors:** Maria Vidal-Sabanés, Raúl Navaridas, Núria Bonifaci, Ada Gay-Rua, Damià Ortega-Peinado, Joaquim Egea, Mario Encinas, Xavier Matias-Guiu, David Llobet-Navas, Xavier Dolcet

## Abstract

High-risk endometrial cancers (EC), such as uterine carcinosarcomas (UCS) and serous endometrial intraepithelial carcinoma (SEIC), are characterized by frequent mutations in tumor suppressor genes (TSGs) and poor clinical outcomes. Traditional genetically engineered mouse models are limited in flexibility and scalability to study the cooperative effects of multiple TSG alterations. Here, we use a multiplexed CRISPR/Cas9-based approach to simultaneously edit the top ten TSGs commonly mutated in high-risk EC directly in the mouse endometrium via intrauterine electroporation. Using rolling circle amplification (RCA) and next-generation sequencing, we demonstrate that this method induces targeted gene editing in a mosaic manner, mimicking tumor heterogeneity. We demonstrate that this approach generates histologically and molecularly faithful models of SEIC and UCS. Importantly, some edited tissues remained histologically normal, emphasizing the complex multistep nature of endometrial tumorigenesis. These CRISPR/Cas9-generated murine models serve as robust platforms to dissect the molecular underpinnings of high-risk endometrial cancer and to accelerate preclinical evaluation of novel therapeutic strategies.

## INTRODUCTION

In the western world, endometrial carcinoma (EC) is the fourth most common cancer among women and the most common of the female genital tract, with an estimated incidence at 10-20 per 100,000 women^1,2^. EC can be classified from pathological, histological or molecular point of view^3^. Classically, EC was classified on basis of clinical and pathological features into two groups: Type I or endometroid endometrial carcinoma (ECC) and Type II or non-endometroid endometrial carcinomas (NECC)^4^. Regarding the histological classification, EC are grouped in endometroid endometrial carcinomas (including its variants), clear cell endometrial carcinomas (CC), serous endometrial carcinomas (SEC) or uterine carcinosarcomas (UCS)^3,5^. The former molecular profiling of EC performed by The Cancer Genome Atlas (TCGA) consortium in 2013 resulted in a new classification of EC in four different groups based on their molecular alterations. These EC groups include: ultramutated, characterized by pathogenic somatic mutations in the exonuclease domain of the replicative DNA polymerase epsilon (*POLE*); hypermutated/microsatellite unstable group, characterized by microsatellite instability as a result of defective mismatch repair (MSI); Copy Number Low (CNL), composed of *P53* wild-type and *POLE* wild-type, MMR-proficient tumors with relatively low somatic copy number alterations and; Copy Number High (CNH)-Serous-like, characterized by mutations in p53 and extensive somatic copy number alterations^6^. Molecular classification provided a relevant advance in the diagnostic, prognostic, and therapeutic management of EC. The vast majority (around 90%) of high-grade non-endometroid ECs, including SEC and UCS, harbor molecular alterations in *p53*^3^. EC containing p53 mutations are considered high-risk cancers that have reliably been associated with the poorest clinical outcomes, high risk of recurrence and particularly high mortality (50-70 % of EC deaths)^5,2^. Mutations in p53 gene are accompanied with highly frequent mutations in other tumor suppressor genes that cooperate with *p53* to derive in high-risk ECs. Histologically, high-risk ECs are mostly classified as UCS, SEC, or CC ECs^3^. UCS are typically biphasic neoplasm harboring carcinomatous and sarcomatous compartments and molecular heterogeneity^7,8^. Mounting molecular and cellular results support that mesenchymal cells of the sarcomatous component are the result of a EMT of epithelial cells of the carcinomatous fraction of CS ^9,10^. SEC usually arise from precursor serous endometrial intraepithelial carcinoma (SEIC) which are an endometrial polyp of atrophic endometrium^11,12^. Histologically, SECs display glandular/papillary architecture, marked cytologic/nuclear atypia and high mitotic index^11,12^. The role of individual or combinations of TSG mutations in the development and progression of EC is still challenging. In fact, oncogenic mutations have been recently found in normal endometria^13^. Therefore, the role of individual TSG and combined TSG mutations in the development, progression and prognostic of EC requires functional validation. Among all TSGs mutated in high-risk EC, some of them are classical and whose function has been broadly analyzed in EC, but others are still not functionally validated in cancer in general or in EC. Moreover, cancers are molecularly heterogeneous, with several populations of cells harboring different sets of molecular alterations co-existing in tumoral parenchyma. Even less known are the clonal collaborative effects of different mutated TSG in EC development. Finally, there is a more than an imperfect correlation between histological molecular and histological classification of ECs. To substantiate such shortcomings, we have used pooled in vivo electroporation of ribonucleoproteins (RNPs) targeting the top ten tumor suppressor genes mutated in high-risk ECs (mainly UCS and serous ECs) to investigate its resulting histological manifestation and aggressiveness.

## RESULTS

### Identification of most frequently mutated tumor suppressor genes in high-risk endometrial cancer

The first comprehensive analysis of endometrial cancers established a molecular classification of EC that correlated prognosis^6^. To model the most aggressive ECs associated with high-risk of recurrence and poor prognosis, we first performed a search from publicly available genomic data (Figure 1A). Candidate TSGs or genes with unknown function in EC retrieved from a search in 8 non-redundant data set studies (total of 1133 samples) available at cBioPortal (https://www.cbioportal.org)^14^. Data from the 8 selected studies was first filtered for mutation profile, histologic grade, and survival data availability (808/1133 samples). Then, samples were further filtered the histologic cancer types associated with high-risk or low risk irrespective of genomic data. High risk selection included the following criteria: high-grade (grade 3) endometroid endometrial carcinoma, uterine serous carcinoma/papillary serous, uterine clear cell carcinoma, uterine carcinosarcoma/uterine malignant mixed mullerian Tumor (334 samples). Low risk included low grade (grades 1 and 2) endometroid ECs (479/808 samples). Uterine undifferentiated, uterine dedifferentiated carcinoma and neuroendocrine carcinomas were excluded because of the low input of samples (3, 1, 1, respectively). High risk selection was confirmed by survival comparison with low-risk samples (Figure 1A). Then, the top ten known TSGs or candidate TSGs after applied high-risk criteria were selected to be targeted by CRISPR/Cas9 edition (Figure 1A and Supplementary Table SR1). For this purpose, we designed crRNA specific to each selected gene, with all crRNA sequences detailed in Supplementary Table SM1. Intrauterine delivery and electroporation of Cas9 ribonucleoproteins (RNPs), as previously described by our laboratory^15^, will produce multiple combinations of gene edits, reflecting the inherent heterogenicity typical of tumor biology (Figure 1A).

**Figure 1.**
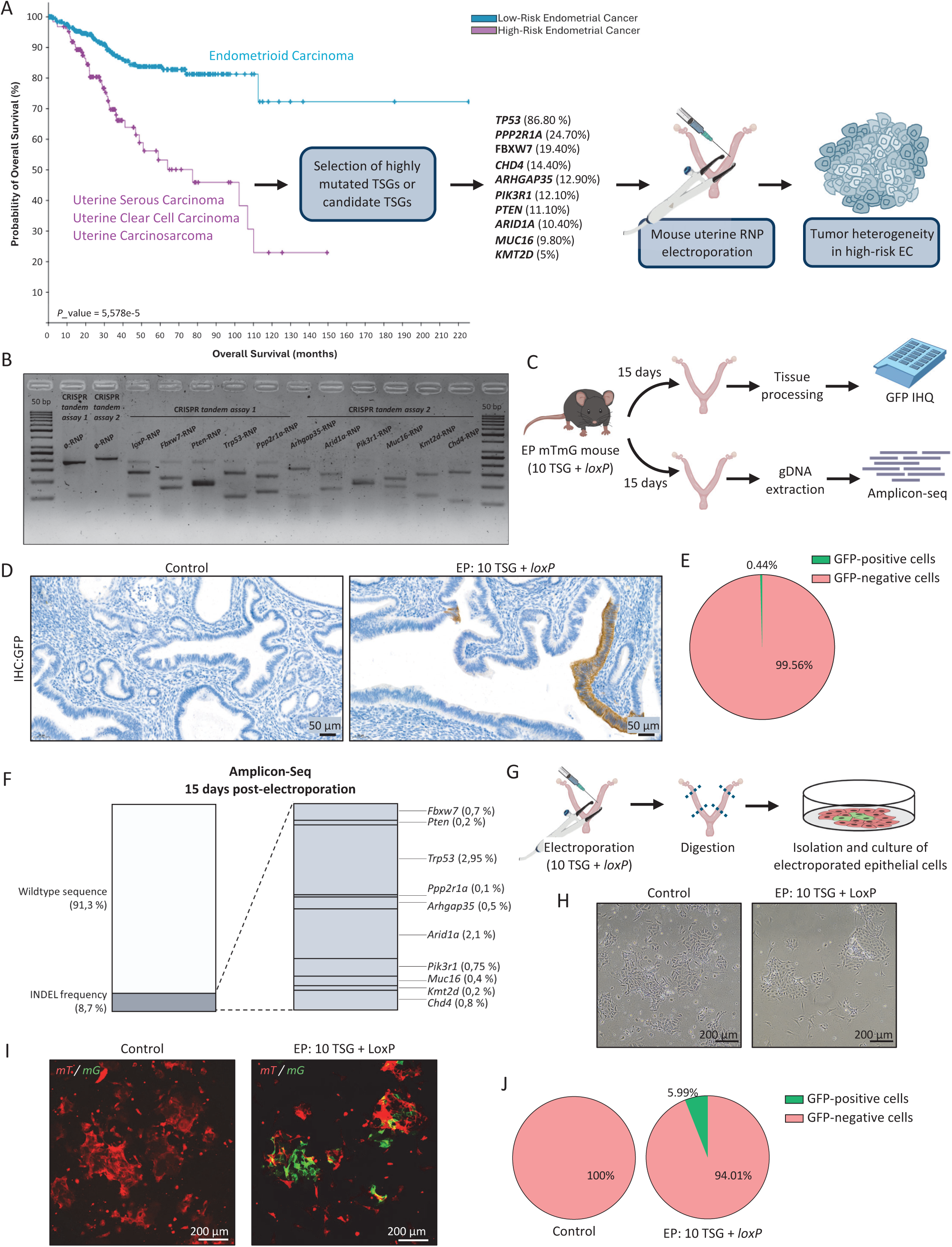
CRISPR/Cas9 editing of frequently mutated tumor suppressor genes in high-risk endometrial cancer. **(A)** Illustration of the experimental design: Kaplan-Meier curve, comparing overall survival from low-risk endometrial cancer subtypes and high-risk endometrial cancer subtypes, obtained from cBioPortal with a *P*-value of 5.578e-5. Most frequently mutated tumor suppressor genes (TSG), or candidates TSG, were selected from high-risk EC group, selected from cBioPortal, and validated in OncoKB. **(B)** Representative image of an agarose-resolved in vitro CRISPR/Cas9 digestion of all RNP target sequences. **(C)** Diagram of the experimental design for validation of in vivo CRISPR/Cas9-mediated gene editing. **(D)** Representative image of GFP immunohistochemistry (IHC) of non-electroporated (control) and electroporated with a pool of RNPs (EP: 10 TSG + *loxP*). Scale bars: 50µm (30X magnification). **(E)** Quantification of GFP-positive cells in samples shown in *(C)*. **(F)** Percentages of edited sequences of CRISPR/Cas9-targeted tumor suppressor genes obtained in Amplicon-Seq analysis. **(G)** Workflow for electroporated endometrial epithelial cells culture. **(H)** Representative phase-contrast images of non-electroporated (control) and electroporated with a pool of RNPs (EP: 10 TSG + *loxP*) epithelial endometrial cells 48h after being cultured. Scale bars: 200µm (20X magnification). **(I)** Representative fluorescence images of non-electroporated (control) and electroporated with a pool of RNPs (EP: 10 TSG + *loxP*) epithelial endometrial cells 48h after being cultured. Scale bars: 200µm (20X magnification). **(J)** Quantification of GFP-positive cells in samples shown in *(I)*.

### CRISPR/Cas9-mediated editing of TSGs associated with high-risk endometrial cancer

Once the target genes were selected and the corresponding cRNA were designed, we assessed the activity of Cas9-RNPs in vitro using a CRISPR/Cas9 assay with oligonucleotides containing all target sequences in tandem (Supplementary Figure S1A). These oligonucleotides were PCR-amplified to generate double-stranded DNA, which was then incubated individually with each RNP. In every case, RNPs efficiently cleaved their respective target sequence (Figure 1B). To assess in vivo activity of RNPs we took advantage of the reporter mouse model expressing an mTmG cassette (Supplementary Figure S1B). Uterine horns of mTmG mice were electroporated with a pooled RNP mix, containing equimolar concentrations, targeting *loxP* and the 10 selected TSGs (*Fbxw7*, *Pten*, *Trp53*, *Ppp2r1a*, *Arhgap35*, *Arid1a*, *Pik3r1*, *Muc16*, *Kmt2d* and *Chd4*). Fifteen days after electroporation, mice were euthanized, and uterine tissues were collected for either GFP immunohistochemistry (IHC) or genomic DNA (gDNA) extraction, followed by amplicon sequencing to detect editing events in the targeted TSGs (Figure 1C). As anticipated, all RNPs individually demonstrated robust activity in vitro, and in the immunohistochemistry assay we observed a small proportion of cells expressing GFP (Figures 1D and 1E). Similarly, as expected by the small fraction of electroporated cells compared to non-transfected cells, amplicon sequencing revealed low but detectable percentages of edited sequences (Figure 1F). The sequences corresponding to the most frequent edits for each gene are presented in Supplementary Table SR2. After detecting edits in the TSGs and *loxP* sites (resulting in GFP expression), we next sought to determine whether electroporated cells could be cultured following electroporation. Following the diagram in Figure 1G, we found that edited epithelial cells were able to grow successfully in vitro, forming viable and healthy cultures (Figure 1H). Furthermore, this approach enabled the quantification of edited cells, at least for *loxP* sites, by analyzing GFP expression (Figures 1I and 1J). These findings demonstrate not only the viability of edited cells in culture, but also the possibility of monitoring and quantifying genome editing efficiency in endometrial-derived cultures following electroporation.

### Multiplexed CRISPR/Cas9-mediated editing of TSGs results in heterogenous populations of epithelial endometrial cells

Amplicon sequencing provides an overall view of CRISPR/Cas9-mediated gene editing. However, it lacks single-cell resolution and cannot determine the specific combination of induced-mutations within individual cells that is the basis of heterogeneity generated by in vivo CRISPR/Cas9 editing^15^. To demonstrate that multiplexed CRISPR/Cas9-mediated editing of TSGs results in mutational heterogeneity, we applied a multiplexed Rolling Circle Amplification (RCA) protocol that allows detection of wildtype TSG mRNAs but not TSG mRNAs containing indels (Figure 2A). First, to demonstrate that all 10 TSG mRNAs were detectable, we performed RCA using padlock targeting individual mRNAs and we observed that all mRNAs were detectable (Figure 2B) and displayed different expression levels (Figure 2E).

**Figure 2.**
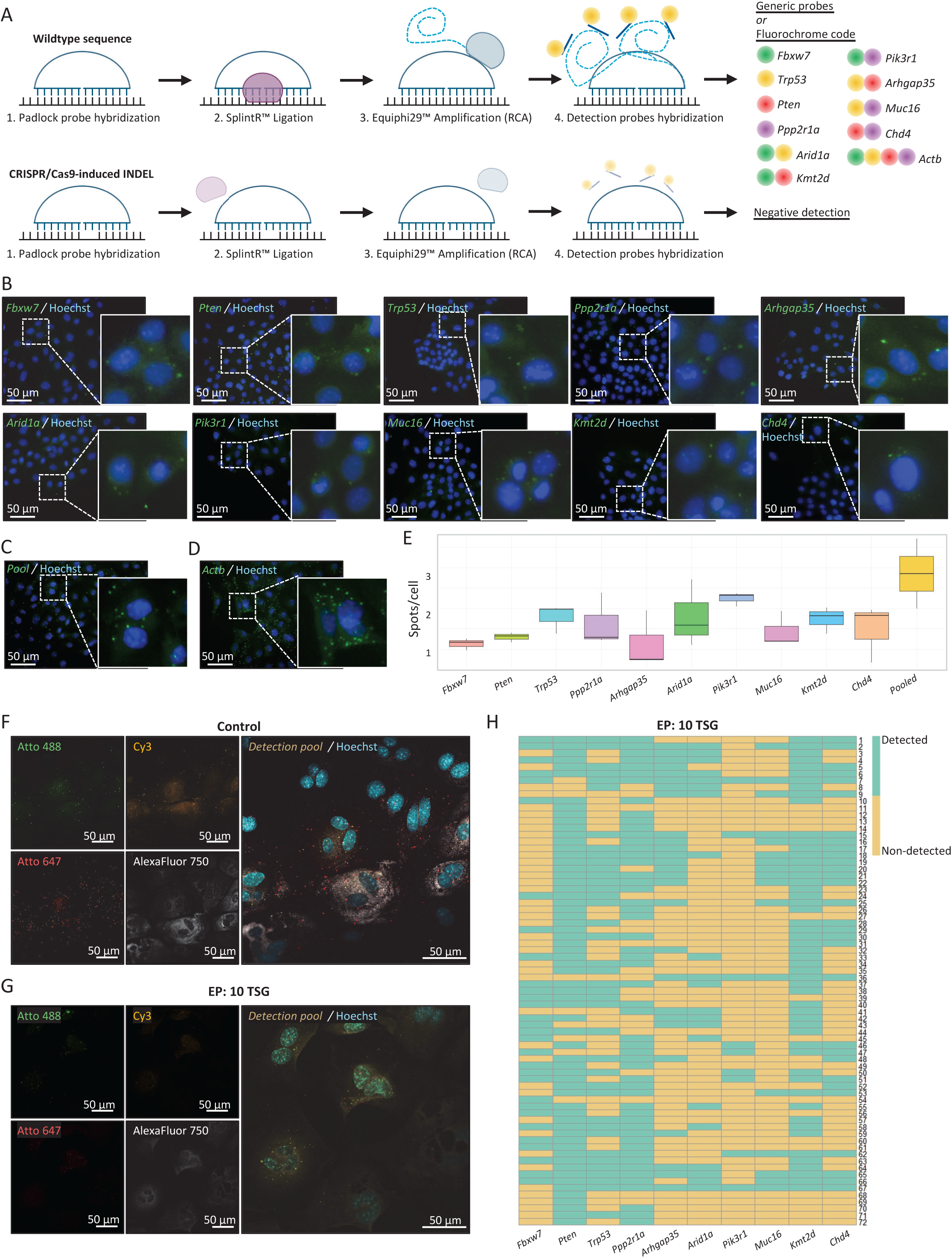
Intrauterine delivery of CRISPR/Cas9 system against most frequently mutated tumor suppressor genes in EC leads to heterogenous populations of epithelial endometrial cells. **(A)** Graphic representation of the spatial mRNA-based method used for validation of tumor heterogeneity generated with CRISPR/Cas9. Briefly, a padlock probe is hybridized in the mRNA of the specific gene, designed at the Cas9-cutting site. Then, SplintR ligase is used to ligate de padlock probe, but if there is not 100% homology between the padlock probe and the mRNA (edited sequence), ligation does not occur. Next step is Rolling Circle Amplification (RCA), where EquiPhi29 polymerase amplifies the padlock sequence. Finally, specific regions of different padlock probes are detected with fluorochrome-labelled probes. If the detection is individual for a single gene, generic probes are used. But if the detection is pooled, a fluorochrome code is used. **(B)** Individual detection of selected TSGs (*Fbxw7, Pten, Trp53, Ppp2r1a, Arhgap35, Arid1a, Pik3r1, Muc16, Kmt2d* and *Chd4*) in wildtype epithelial endometrial cells detected with generic probes (green). Hoechst (blue) is used for nuclear staining. Scale bars: 50µm (40X magnification). **(C)** Pooled detection of mRNA from selected TSGs in wildtype epithelial endometrial cells detected with generic probes (green). Hoechst (blue) is used for nuclear staining. Scale bar: 50µm (40X magnification). **(D)** Individual detection of *Actb* mRNA with generic probes. Scale bar: 50µm (40X magnification). **(E)** Quantification of individual and pooled detections of selected TSG in wildtype cells, shown in *(B)* and *(C)*. **(F)** Representative image of pooled detection of selected TSG in non-electroporated (control) epithelial endometrial cells, detected with fluorochrome code. Images show detection with Atto488 (green), Cy3 (orange), Atto647 (red) and/or AlexaFluor750 (grey). Hoechst (blue) is used for nuclear staining. Scale bars: 50µm (60X magnification). **(G)** Representative image of pooled detection of selected TSG in electroporated with a pool of RNPs (EP: 10 TSG), detected with fluorochrome code. Images show detection with Atto488 (green), Cy3 (orange), Atto647 (red) and/or AlexaFluor750 (grey). Hoechst (blue) is used for nuclear staining. Scale bars: 50µm (60X magnification). **(H)** Quantification and co-localization of RNA spots according to the fluorochrome code at a single cell resolution. Binary heatmap represents the presence (green) or absence (yellow) of *Fbxw7, Pten, Trp53, Ppp2r1a, Arhgap35, Arid1a, Pik3r1, Muc16, Kmt2d* or *Chd4* detection in each cell (rows).

Once demonstrated that all padlocks were suitable for detection of individual mRNAs, we proceeded to multiplexed RCA analysis using a pool of all padlocks that were detected by probes with a fluorochrome code, as depicted in Figure 2A and Supplementary Figure S2A. To prove that simultaneous detection of all fluorochromes, used in detection probes for targeted genes, was possible, we performed RCA to Actin B mRNA, which is highly expressed in all cells. Detection probe for Actin B mRNA was designed to hybridize with all detection probes conjugated with the four different fluorochromes (Supplementary Figure S2B). To demonstrate the suitability of the method to detect loss of expression of the analyzed mRNA, we performed RCA to analyze *Pten* expression on cells isolated from Cre:ER^T-/-^ *Pten*^F/F^ (Wt) mice or Cre:ER^T+/-^ *Pten*^F/F^ (Pten KO) mice. Only Wt but not Pten KO cells displayed spots corresponding to positive RCA staining (Supplementary Figure S2C). Next, we performed RCA for detection of the mRNA corresponding to the 10 CRIPR/Cas9 targeted genes in endometrial cells isolated from Control (electroporated with no RNP) (Figure 2F) or endometrial cells isolated from endometrium electroporated with the cocktail of RNPs targeting the 10 TSGs (Figure 2G). Spots with fluorochrome codes corresponding to each TSG were quantified in individual cells as described in Material and Methods section. Presence/absence of spots corresponding to each mRNA was highly heterogeneous among the analyzed cells, demonstrating that multiplexed electroporation of RNPs generated high degree of heterogeneity in gene editing across different cells (Figure 2H).

### Multiplexed CRISPR/Cas9-mediated editing of TSGs results in two main types of endometrial lesions

To study the functional consequences of high-risk TSGs in the development of EC, 14 mice were electroporated with pooled RNPs. The health appearance of electroporated mice was monitored weekly and they were sacrificed when signs of pathological condition such as hunched back, loss of weight or motility or vaginal bleeding were observed. Dates of birth, electroporation and sacrifice, weeks of progression and histopathology of endometrial lesions are summarized in Figure 3A and Supplementary Table SR3. Interestingly, 10 of 14 electroporated mice displayed evident macroscopic endometrial lesions, with two different macroscopic appearance (Figure 3B (II) and (III)), the remaining uteri showed normal macroscopic appearance (Figure 3B (I)). Histopathological evaluation diagnosed EC lesions of SEIC or UCS (Figure 3C), that were associated with the two different macroscopic forms (Figure 3D). The percentage of mice displaying normal, SEIC or UCS, is summarized in Figure 3E.

**Figure 3.**
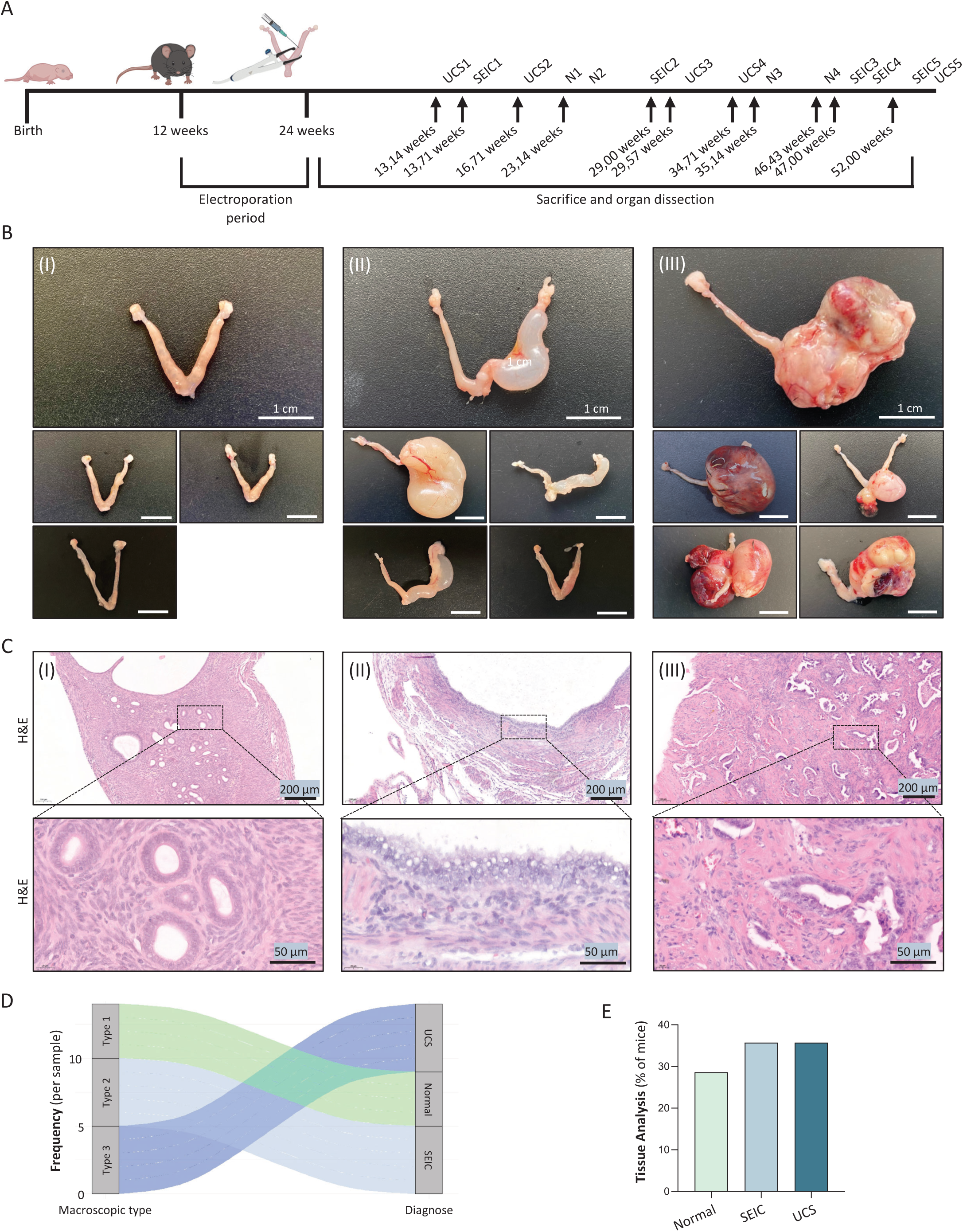
Intrauterine deletion of tumor suppressor genes leads to the formation of high-risk endometrial cancer subtypes. **(A)** Timeline for the *in vivo* analysis of endometrial lesions in electroporated mice. **(B)** Macroscopic images from three different uterine lesions from type 1 (I), type 2 (II) and type 3 (III). Scale bar: 1cm. **(C)** Representative hematoxylin and eosin (H&E) staining of lesion groups (I), (II) and (III) from endometrial sections of EP mice. Scale bars: 200µm (10X magnification) and 50µm (40X magnification). **(D)** Histopathological analysis of endometrial sections shown in *(C)*. Group (I) lesions correspond to normal (N) histology while groups (II) and (III) correspond to Serous Endometrial Intraepithelial Carcinoma (SEIC) and Uterine Carcinosarcoma (UCS), respectively. **(E)** Percentages of each histological subtype.

### Histologically normal endometria display mutations in targeted TSG

Approximately one-third of electroporated mice displayed normal histological morphology, with no signs of cancerous or precancerous lesions (Figure 4A). Therefore, we wondered whether this was caused by a failed electroporation that resulted in the lack of RNP delivery into the endometrial cells. Harnessing the mTmG reporter system, we first performed GFP IHC to detect the switch from mT to mG caused by LoxP targeting RNPs as a readout that electroporation was successfully performed. We found heterogenous positive staining for GFP (Figure 4B), suggesting a successful electroporation and RNP delivery into the cells. Considering the heterogeneity of pooled RNP delivery into the cells^15^, we found unlikely that no other RNP targeting any of the 10 TSG had entered into the cells. To demonstrate the mutational status of the targeted TSGs, we submitted DNA from histologically normal endometrium (N4) to amplicon sequencing. Five out of the 10 targeted TSGs displayed indels (Figure 4C), being *Pten* and *Trp53* the TSGs with a higher percentage of mutated sequences. Among the 5 TSGs with sequence edits, *Pten*, *Trp53*, *Pik3r1* and *Kmt2d* displayed frameshift mutations and *Arhgap35* in-frame mutations (Figure 4D and Supplementary Table SR4).

**Figure 4.**
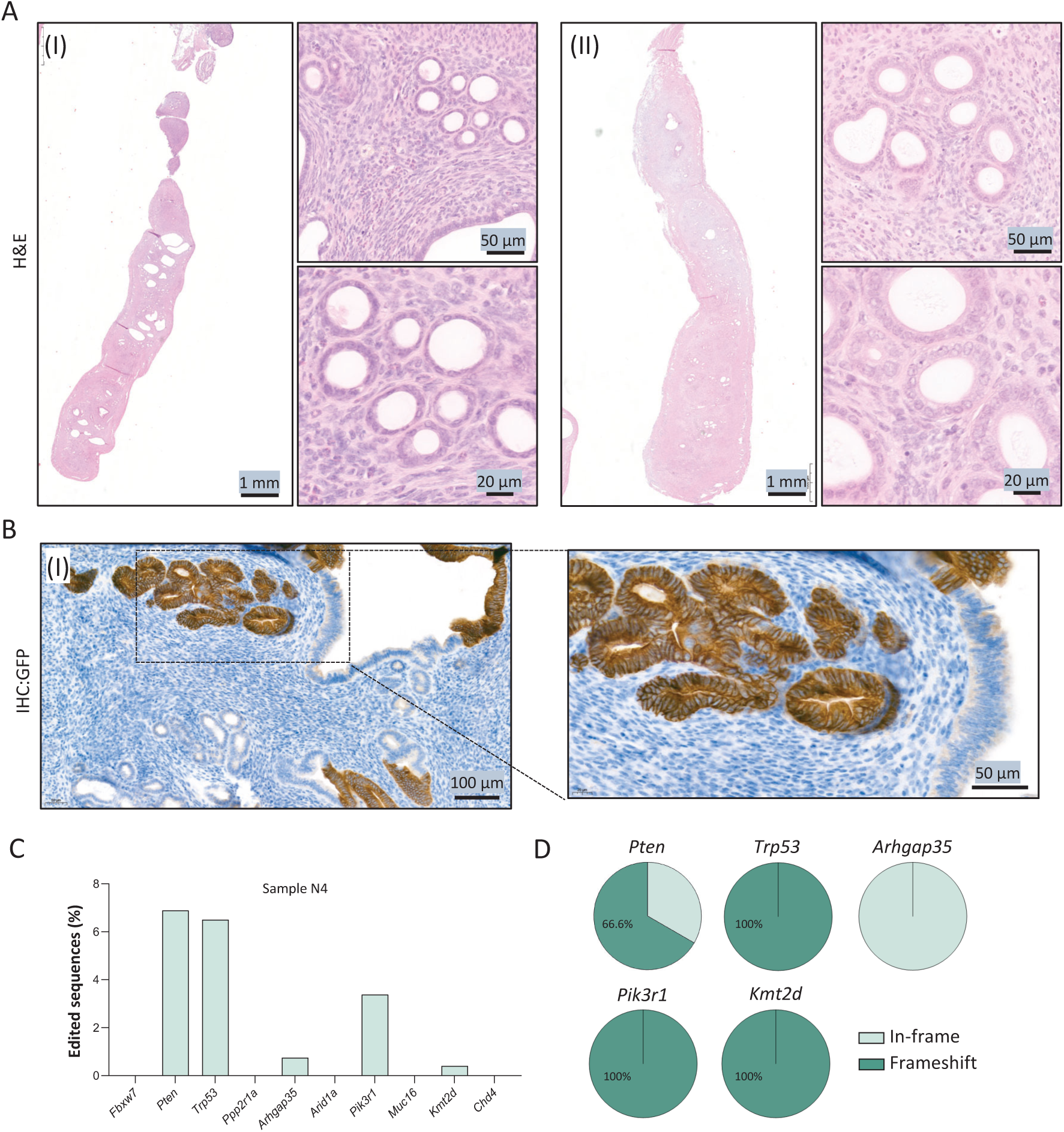
Histologically normal endometrium harbor mutations in targeted genes. **(A)** Hematoxylin and eosin (H&E) staining corresponding to the normal endometria N1 (I) and N4(II). Scale bars: 1mm (1.5X magnification), 50µm (30X magnification) and 20µm (60X magnification). **(B)** Representative images of GFP immunohistochemistry (IHC) analysis in normal type endometrial sections. Scale bars: 100µm (15X magnification) and 50µm (40X magnification). **(C)** Frequency of edited sequences in sample N4 (normal type lesion), showed in %. **(D)** Pie charts showing the proportion of mutation type in edited genes from normal type sample N4.

### UCS and SEIC generated with CRISPR/Cas9 intrauterine delivery exhibit morphological and molecular features of human pathology

Next, we proceeded to a morphological and molecular characterization of SEIC generated by CRSPR/Cas9 system. Accurate histopathological analysis of hematoxylin and eosin (H&E) stained sections of SEIC highlighted typical morphological features of serous pathology^16^ including nuclear atypia, loss of cell polarity, epithelial stratification, presence of epithelial papillary structures and increased numbers of mitotic and apoptotic cells (Figure 5A). One of the hallmarks of SEIC is the presence of *Tp53* mutations. In humans, *Tp53* mutations are usually assessed by immunohistochemistry showing the typical ‘all or nothing’ pattern (strong positivity or total absence of staining), which contrasts with the ‘wildtype’ pattern, characterized by focal, weak, and somewhat variable staining from area to area^16^. IHC analysis of P53 in CRISPR/Cas9-generated SEICs revealed the presence of null and nuclear patterns like those observed in human tissues (Figure 5B, 5C and Supplementary Figure S3A). Interestingly, the only SEIC that was not edited by CRISPR/Cas9 RNP (SEIC4) displayed strong nuclear staining for P53, suggesting that secondary molecular alterations in this gene occurred (Supplementary Figure S3B). Another marker used in differential diagnosis of SEIC is the expression of p16, which is widely increased in serous carcinomas^17^. Consistently, endometria diagnosed of SEIC displayed strong cytoplasmic p16 immunostaining, compared to wild type endometria (Fig 5D, 5E and Supplementary Figure S3C).

**Figure 5.**
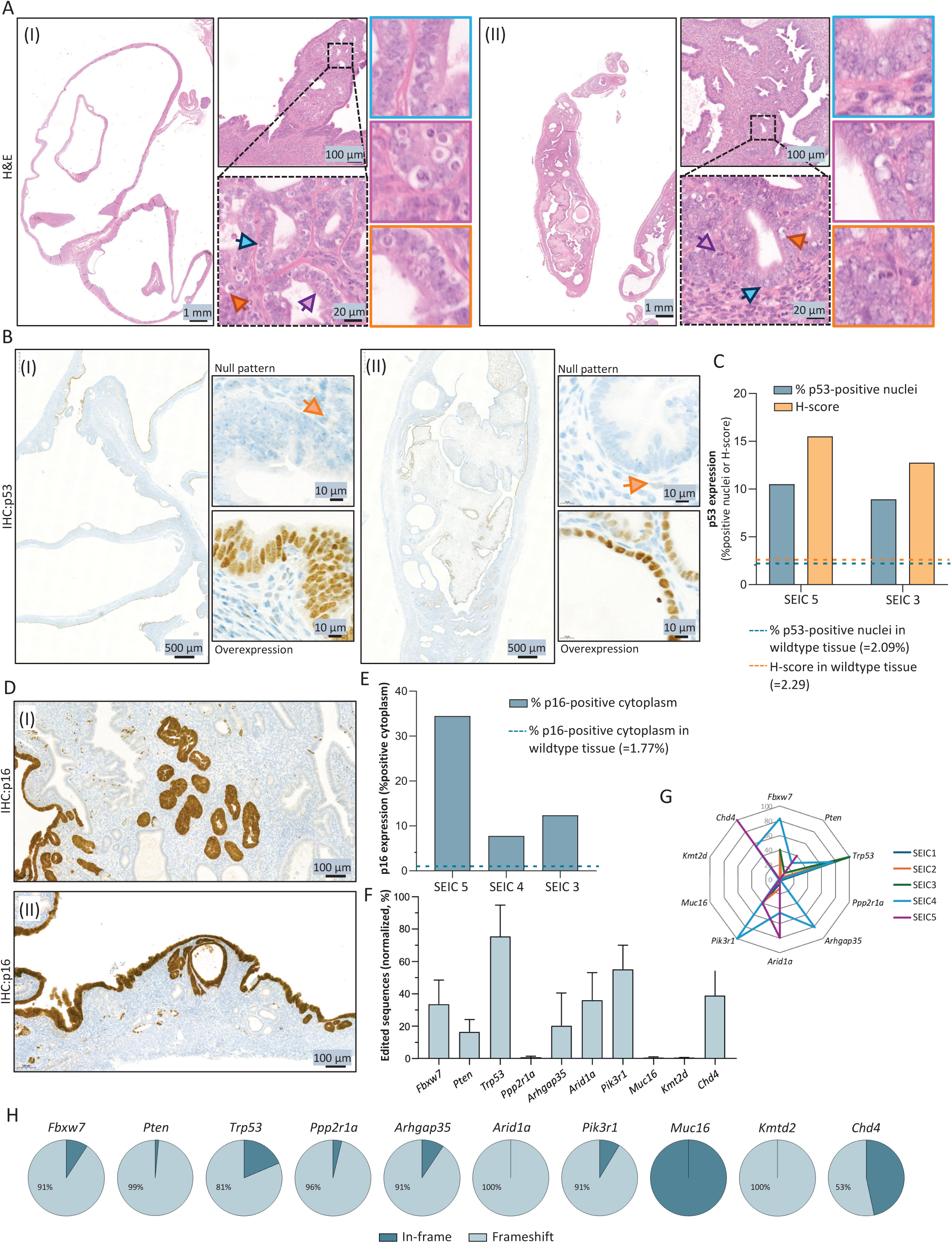
Serous Endometrial Intraepithelial Carcinomas generated with CRISPR/Cas9 intrauterine delivery exhibit morphologic and molecular features of human pathology. **(A)** Hematoxylin and eosin (H&E) staining of Serous Endometrial Intraepithelial Carcinomas (SEIC) cases SEIC3 (I) and SEIC5 (II), highlighting typical morphological characteristics in serous pathology. Blue arrows and squares show pleomorphic nuclei, arrows and squares in purple show apoptotic cells and orange arrows and squares show mitotic cells. Scale bars: 1mm (0.9X magnification), 100µm (10X magnification) and 20µm (60X magnification). **(B)** Immunohistochemistry (IHC) images of p53 in cases SEIC3 (I) and SEIC5 (II), underlining aberrant p53 expression as null pattern (upper ampliation, orange arrow indicates p53 internal positive control) and overexpression pattern (lower ampliation). Scale bars: 500µm (2X magnification) and 10µm (100X magnification). **(C)** Quantification of p53-positive nuclei and p53 histoscore (H-score) in cases SEIC5 and SEIC3. Discontinuous horizontal lines represent % of p53-positive nuclei or p53 H-score in a wildtype endometrium. **(D)** Immunohistochemistry (IHC) images of p16 in cases SEIC3 (I) and SEIC4 (II). **(E)** Quantification of p16-positive cells in cases SEIC5, SEIC4 and SEIC3. Discontinuous horizontal lines represent the % of p16-positive cells in a wildtype endometrium. **(F)** Normalized frequency of edition of selected tumor suppressor genes (*Fbxw7*, *Pten*, *Trp53*, *Ppp2r1a*, *Arhgap35*, *Arid1a*, *Pik3r1*, *Muc16*, *Kmt2d* and *Chd4*) from SEIC cases (SEIC 1-5). **(G)** Spider plot showing the normalized edition frequences of targeted TSG across SEIC samples (SEIC 1-5). Each axis represents one gene (*Fbxw7*, *Pten*, *Trp53*, *Ppp2r1a*, *Arhgap35*, *Arid1a*, *Pik3r1*, *Muc16*, *Kmt2d* and *Chd4*) and the radial distance corresponds to the normalized edition frequency. **(H)** Pie charts showing the proportion of mutation type in edited genes from SEIC samples (SEIC 1-5). Data is represented as the mean of in-frame/frameshift mutations proportion from all samples.

Next, we aimed to investigate the mutational landscape of targeted TSGs in SEIC. For this purpose, DNA extracted from endometria of the 5 mice diagnosed as SEIC (SEIC1-5) was extracted and submitted to NGS-amplicon sequencing. Global analysis of sequences demonstrated the presence of indels in all 10 targeted TSGs with different fractions of edition among the 5 SEIC samples (Figure 5F). Normalized edition frequences of targeted TSG across SEIC samples (SEIC1-5) show heterogeneous percentages of indels among the 10 targeted TSGs (Figure 5G). Noteworthy *Trp53*, which is the most frequently mutated TSG in SEIC, was the one showing the highest percentage of edition. Most frequent indels of SEIC 1 to 5 are shown in Supplementary tables SR5 to SR9. All genes but *Muc16* displayed indels that resulted in frameshift mutations disrupting the open reading frames of the targeted TSGs (Figure 5H and Supplementary Table SR10).

UCS are highly heterogeneous neoplasia characterized by the presence of carcinomatous and sarcomatous compartments^7,8^. Histopathological evaluation of UCS arose from electroporated endometria clearly displayed both sarcomatous and carcinomatous regions (Figure 6A), that were confirmed by immunohistochemistry analysis of vimentin as marker of mesenchymal cells (Figure 6B) and E-Cadherin (Figure 6C) / Cytokeratine-8 (Figure 6D) as markers of epithelial cells. All UCS displayed myometrial invasion and one of them (UCS5) established metastatic outgrowth in the lymph nodes (Figure 6E), that were identified by cytokerain-8 immunostaining (Figure 6F).

**Figure 6.**
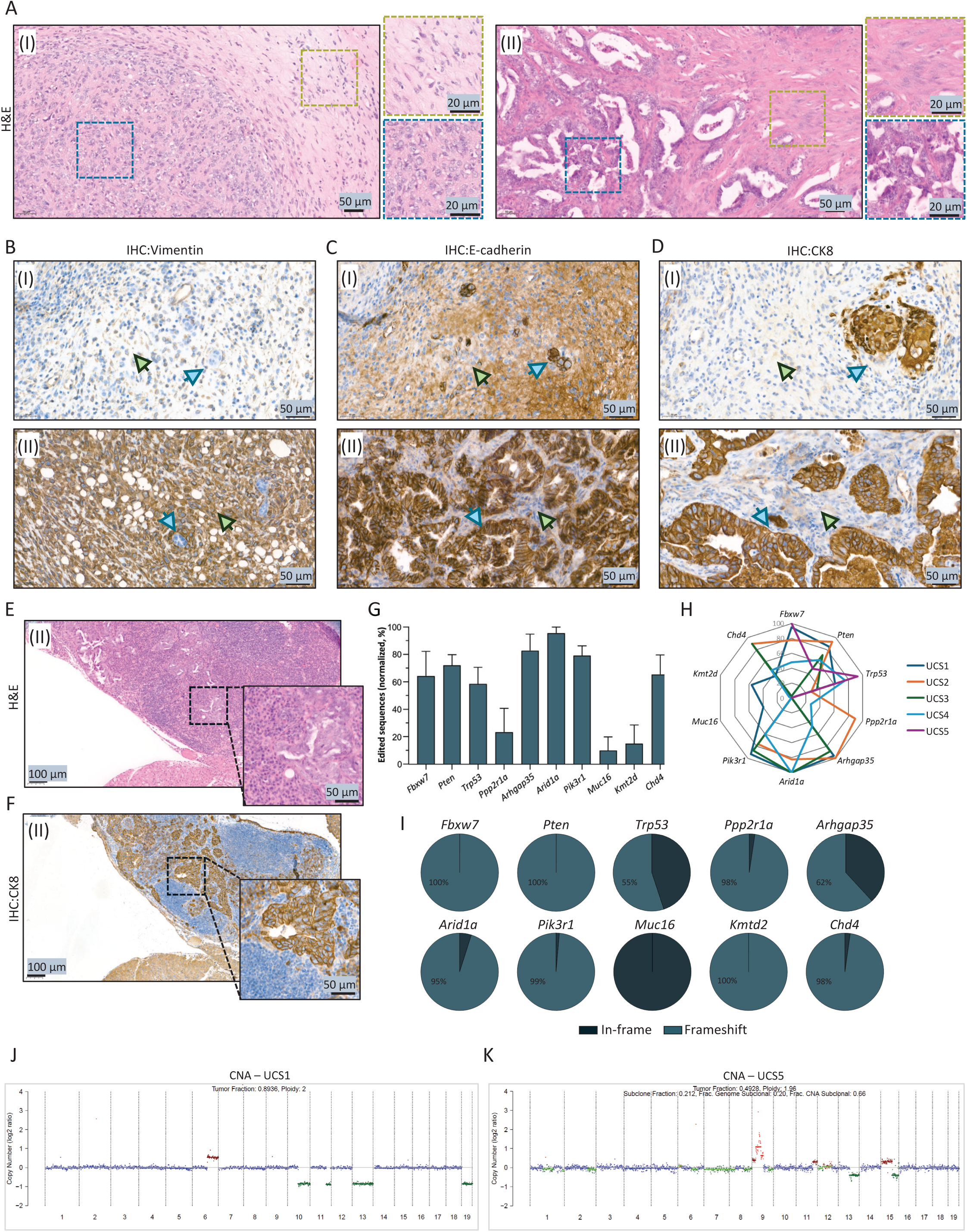
Uterine carcinosarcomas obtained from multiplexed CRISPR/Cas9 display morphologic and molecular characteristics of human disease. **(A)** Hematoxylin and eosin (H&E) staining of Uterine Carcinosarcomas (UCS) cases UCS 2 (I) and UCS 5 (II). Squares in blue indicate epithelial compartment and squares in green show mesenchymal compartment. Scale bars: 50µm (20X magnification) and 20µm (60X magnification). **(B-D)** Representative immunohistochemistry (IHC) images of Vimentin, E-cadherin and Cytokeratin 8 (CK8) detections in cases UCS 2 (I) and UCS 5 (II). Blue arrows indicate epithelial compartment, and green arrows indicate mesenchymal compartments. Scale bar: 50µm (30X magnification). **(E)** Hematoxylin and eosin staining (H&E) of lymph node metastasis in case UCS 5 (II). Scale bars: 100µm (10X magnification) and 50µm (40X magnification). **(F)** Immunohistochemistry (IHC) of Cytokeratin 8 (CK8) of lymph node metastasis in case UCS 5 (II). Scale bars: 100µm (10X magnification) and 50µm (40X magnification). **(G)** Normalized frequency of edition of selected tumor suppressor genes (*Fbxw7, Pten, Trp53, Ppp2r1a, Arhgap35, Arid1a, Pik3r1, Muc16, Kmt2d* and *Chd4*) from UCS cases (UCS 1-5). Data is represented as mean ± S.E.M. **(H)** Spider plot showing the normalized edition frequences of targeted TSG across SEIC samples (SEIC 1-5). Each axis represents one gene (*Fbxw7, Pten, Trp53, Ppp2r1a, Arhgap35, Arid1a, Pik3r1, Muc16, Kmt2d* and *Chd4*) and the radial distance corresponds to the normalized edition frequency. **(I)** Pie charts showing the proportion of mutation type in edited genes from UCS samples (UCS 1-5). Data is represented as the mean of in-frame/frameshift mutations proportion from all samples. **(J)** Copy number variations of UCS1. **(K)** Copy variation analysis of UCS5. Copy number alterations (CNAs) were estimated using the ichorCNA algorithm. The X-axis represents genomic coordinates concatenated by chromosome (1–22), and the Y-axis displays the logz copy number ratio for each genomic bin. Each data point is color-coded according to the estimated integer copy number: 1 copy = dark green, 2 copies = blue, 3 copies = brown, ≥4 copies = red. Horizontal lines represent the segment median values, with color matching the corresponding copy number state. Segments are colored only if the event is predicted to be clonal; subclonal events are not annotated with segment lines. These profiles reflect the expected resolution of shallow whole-genome sequencing and ichorCNA’s probabilistic segmentation model (10.1038/s41467-017-00965-y, 10.1158/1078-0432.CCR-21-2291).

DNA extracted from endometria of the 5 mice diagnosed as UCS (UCS1-5) was also submitted to NGS-amplicon sequencing to identify the TSGs that suffered indels. Like SEIC, normalized editing frequences of the targeted TSG in all UCS samples (UCS1-5) demonstrated that all genes were targeted across the different samples (Figure 6G), but the percentages of indels varied heterogeneously for each TSG within individual samples (Figure 6H). Most frequent indels of UCS1 to 5 are shown in Supplementary tables SR11 to SR15. All genes but *Muc16* displayed indels that resulted in frameshift mutations disrupting the open reading frames of the targeted TSGs (Figure 6I and Supplementary Table SR16).

### UCS acquired high copy number variations

Integrated genomic characterization of EC clustered the vast majority of UCS to the CNV-H molecular subgroup of EC^3,6^. Therefore, we questioned whether CRISPR/Cas9 introduced mutations in the 10 selected genes resulted in increased copy number variations. To assess chromosomal instability across tumors, we performed shallow whole-genome sequencing (sWGS) on two primary endometrial cancer samples and inferred copy number alterations (CNAs) using the ichorCNA algorithm (10.1038/s41467-017-00965-y). The resulting genome-wide profiles revealed striking differences in CNA burden and clonal architecture between the two tumors.

UCS1 exhibited a largely copy-number neutral genome, with the majority of genomic bins showing a logz ratio near zero and an estimated tumor fraction of 0.89 and ploidy of 2.0. Only a few focal CNAs were detected, including heterozygous deletions on chromosomes 10, 11, 13, and 19 (1-copy losses), and a single low-level gain on chromosome 6 (3 copies). These alterations were largely clonal, as reflected by the presence of corresponding segment medians. The overall CNA pattern is consistent with a genomically stable tumor harboring limited chromosomal aberrations (Figure 6J and Supplementary Table SR17). In contrast, UCS5 displayed extensive genomic alterations despite a near-diploid ploidy (1.96). This tumor exhibited a markedly higher CNA burden, with multiple segmental gains and high-level amplifications (≥4 copies) across chromosomes 6, 9, 11, and 12, alongside widespread heterozygous deletions (1-copy losses) throughout the genome. The estimated tumor fraction was 0.49, and the copy number profile revealed substantial subclonality, with 20% of the genome and 66% of CNA segments predicted to be subclonal (Figure 6K and Supplementary Table SR17). These features suggest ongoing chromosomal instability and intratumoral heterogeneity, despite the near-diploid ploidy estimate.

Together, these results highlight the heterogeneity in genomic architecture among primary tumors, with implications for tumor evolution, clonal dynamics, and potential therapeutic vulnerabilities.

## DISCUSSION

In the current work, we have used CRISPR/Cas9 technology to introduce mutations to TSG associated with high-risk ECs. Interestingly, we have found that multiplexed editing of the 10 most frequently mutated TSG in such ECs recapitulates histological manifestations associated with high-risk ECs, such as UCS and SEIC.

Genomic data provided by seminal TCGA study and later GWS studies, retrieved lists of already known TSG mutations but also identified new mutations. For instance, these studies confirmed the presence of well-known tumor suppressor genes in EC such as *Pten* or *Tp53* but also unveiled the presence of mutations in TSG that had not previously been found to be altered in EC and identified several new molecular alterations in genes with unknown functions. Many of these newly identified alterations are of unknown significance and their role as a putative driver mutation remains to be ascertained. Therefore, it is crucial to directly assess the functional contribution of these alterations to the tumor phenotype. This approach enables the analysis of the roles of single mutations or mutation combinations on tumor phenotypes, and may facilitate the analysis of tumor behavior and may uncover new therapeutic targets for highly aggressive forms of EC.

As most malignancies, EC arises from the accumulation of driver mutations that confer the hallmarks of cancer cell phenotype^33^. Whole genome sequencing have unveiled or confirmed previously known mutations in EC^6^. However, functional validation of the identified alterations as a driver mutation itself or in combination with other alterations is still challenging in cancer research. Traditionally, generation of either constitutive or conditional genetically modified mouse models has been the main approach to interrogate the function of candidate TSG in the development of EC. Such approaches provided unfading information about the role of genes alterations in EC development and progression. However, one of the main bottlenecks of this approach is the study of collaborative, additional or synergic roles of TSGs in EC. The generation of mouse models carrying endometrial-specific mutations in more than one TSG (i.e., double or triple knock-out), which require breeding several TSG conditional mouse strains with appropriate mouse carrying tissue specific Cre recombinase, makes nearly impossible to study the effects of multiple mutations at same time. Moreover, many conditional genetically modified mouse models recapitulate endometroid EC type but there is a very limited availability of mouse models developing high-risk EC such as serous, clear cell or carcinosarcomas^34,35^. To date, there are very limited genetically modified mouse models to model SEIC or SEC. The development of mouse models of SEIC or SEC is specially challenging. Mice with endometrium-specific deletion of *Trp53* developed type II EC including serous, clear cell, undifferentiated EC and UCS^36^. Conversely, other studies using conditional p53 knock-out crossed with mice lacking other TSGs have shown that loss of p53 alone is not sufficient to cause EC. Conditional deletion of Pten/p53^37^, Cdh1/p53^38^, Pot1a/p53^39^, or Rb1/p53^40^ in mouse endometrium resulted in EC but, in all these studies, control mice lacking only p53 had no signs or low incidence and penetrance of EC. Another study demonstrated that conditional loss of p53 in the presence of PIK3CA^H1047R^ mutation drives hyperplasia and endometrial intraepithelial carcinoma, but control mice lacking only p53 had no endometrial lesions^37^. Recent studies developed mouse models of SEC generated by orthotopic transplantation of CRISPR/Cas9 edited mouse organoids^41^. Our recently described protocol for in vivo CRISPR/Cas9 editing of endometrial epithelial opened a new door to future rapid, flexible, customizable and multiplexable in vivo modeling of EC^42^. Using this approach, we have recently demonstrated that concomitant deletion of *Pten* and *Trp53* definitely leads to development of UCS^10^, but in this previous study we did not identify any case of SEIC or SEC.

The results from amplicon sequencing from pooled multiplexed RNP electroporation demonstrated that in vivo editing cause mutations in all 10 targeted TSGs with one single electroporation. Noteworthy, we demonstrated that only a small fraction of cells electroporated compared to non-transfected cells and amplicon sequencing revealed low, but detectable percentages of edited sequences. This feature may provide better modeling of cancer origin over the classical tissue specific knock-out. Most of conditional Cre-*LoxP* systems used to knock-out genes involved in EC, cause the ablation of the floxed gene in all cells expressing the Cre recombinase. Even Cre expressing mouse models that allow specific deletion of genes in endometrial epithelial cells genes deletion takes place in all epithelial cells, leaving no wildtype epithelial endometrial cells in the whole uterus. In contrast, CRISPR/Cas9 mediated edition of mouse endometrial epithelial cells takes place in individual cells, resulting in a mosaic of edited cells side-by-side with wild type cells, which provides a closer scenario to tumoral microenvironment composition. Moreover, multiplexed electroporation of CRISPR/Cas9 RNP targeting different genes generates a high degree of molecular heterogeneity. As we have demonstrated by in situ analysis of targeted TSGs RNA at single cell level, electroporation leads to molecular heterogenous populations of cells. It is easy to speculate that cells receiving most advantageous combinations of mutations will be the ones that participate in tumor development.

One of the most surprising results was the presence of indels in histologically normal endometria. Most of these indels caused frameshift mutations that resulted in open reading frame alterations. It has been demonstrated that normal human endometria harbor oncogenic driver mutations in more than one gene^13,43^. This phenomena is not restricted to the endometria and age related increase of mutations in cancer driver genes has been reported in many tissues^44–47^. Therefore, our findings provide a functional demonstration that the induction of oncogenic mutations itself is not sufficient for EC development.

One of the most remarkable findings of our study is that the histological outcomes resulting from high-risk associated TSG editing align closely with data obtained from GWS of human ECs. Noteworthy, introduction of high-risk associated mutations recapitulates histological features of human EC. None of the mice electroporated with TSG-targeting RNPs pool resulted in the development of low-grade endometroid carcinomas, which are low risk ECs associated with good prognosis. Instead, all ECs were classified as UCS or SEIC, two histological types strongly associated with highly aggressive ECs.

UCS characterizes one of the most extreme illustrations of tumor heterogeneity among human cancers^7^. Most UCS belong to the copy-number high serous-like molecular subtype of endometrial carcinoma, are frequently accompanied by many gene copy-number alterations. Here, we analyzed CNV in two UCS and we found that one of them (UCS5) presented high CNV, while the other one was largely copy-number neutral genome (UCS1). This can be explained because a proportion of cases (20%) probably represent the progression of tumors initially belonging to the copy-number low endometrioid-like molecular subtype^7^. Supporting this hypothesis, UCS1 had a progression of 13-14 weeks whilst UCS5 was the result of 52 weeks of progression after electroporation. Moreover, previous studies demonstrated a subset of UCS belong to the CNV-L EC molecular classification^48^. Besides the biological significance, CNV may have prognostic and therapeutic value^49^.

Although non-invasive lesion, SEIC is considered a high-risk endometrial lesion that is the precursor of SEC^50^. At the molecular level, both SEIC and SEC are characterized by *Trp53* mutations. Consistently with this data, 4 out of 5 SEIC developed after electroporation displayed CRISPR/Cas9 edits in *Trp53* gene, and the remaining SEIC, although non-targeted by the CRISPR/Cas9, presented zones with strong nuclear P53 staining, suggesting secondary mutations in *Trp53* during tumor progression. Unlike other uterine precursor lesions, diagnostic and management strategies for SEIC remain varied. There are limited guidelines available for the surgical management of SEIC and significant heterogeneity in management strategies^51^. The lack of current models for SEIC or SEC boosts the potential use of these models as a platform for better understanding its molecular biology and the assessment of potential therapeutic interventions.

In summary, our study provides compelling evidence that multiplexed CRISPR/Cas9-mediated editing of tumor suppressor genes in the mouse endometrium effectively models the molecular, morphological, and genomic hallmarks of high-risk endometrial cancers. By recapitulating the heterogeneity, progression, and histological diversity of human UCS and SEIC, this approach bridges a critical gap in the functional validation of cancer mutations. Moreover, the presence of oncogenic mutations in histologically normal endometria emphasizes the complex multistep nature of endometrial tumorigenesis. This platform holds significant promise for dissecting the cooperative roles of cancer genes, testing novel therapeutic strategies, and refining our understanding of tumor evolution in aggressive gynecological malignancies.

## MATERIAL AND METHODS

### Experimental mouse models

Mice were housed in a barrier facility, and pathogen-free procedures were used in all mouse rooms. Animals were under 12 hours of light/dark cycles at 22°C, and they had ad libitum access to water and food. All procedures were performed according to the guidelines of the Ethical Committee of Universitat de Lleida and the National Institute of Health Guide for the Care and Use of Laboratory Animals. Homozygous membrane-targeted tandem dimer Tomato/membrane-targeted Green Fluorescent Protein (mTmG) reporter (B6.129(Cg)-Gt(ROSA)^26Sortm4^(ACTB–tdTomato,–EGFP)^Luo^/J) were obtained from the Jackson Laboratory (Bar Harbor, ME, USA). Wild-type C57BL/6J were bred in Universitat de Lleida animal housing facility. Primers and PCR conditions for genotyping are detailed in Supplementary Table SM1.

### Preparation of CRISPR/Cas9-ribonucleoprotein

RNP preparation was performed as previously described^15^ with some modifications. CRISPR RNAs (crRNAs) and trans-activating CRISPR RNAs (tracrRNA) were obtained from Integrated DNA Technologies (IDT). Sequences of crRNA used for this work are detailed in supplementary material (Supplementary Table SM2). Lyophilized crRNAs and tracrRNA were resuspended at 100µM in nuclease-free duplex buffer, containing 100mM potassium acetate and 30mM HEPES, pH 7.5, (11-05-01-03, IDT). For crRNA:tracrRNA hybridization, both were mixed at equimolar concentration and diluted in nuclease-free duplex buffer to a 20µM concentration. Then, RNAs were heated at 95°C for 5 minutes and subjected to a negative temperature gradient (-2°C/30sec) in a thermal cycler until they reached room temperature. Hybridized 20µM crRNA:tracrRNA (single guide RNA, sgRNA) stocks were stored at -20°C. Recombinant Streptococcus pyogenes Cas9 (Alt-R® S.p. Cas9 Nuclease V3, 1081059, IDT) was mixed at equimolar concentration with sgRNA at a final concentration of 15µM in 4µl (0.5µl of 61µM Cas9 and 1.5µl of 20µM sgRNA). The mixture was incubated for 20 minutes at room temperature to allow Cas9-sgRNA RNP formation.

### In vitro CRISPR/Cas9 ribonucleoprotein assay

To test the Cas9 nuclease on-target activity of *loxP*-RNP, *Fbxw7*-RNP, *Pten*-RNP, *Trp53*-RNP, *Ppp2r1a*-RNP, *Arhgap35*-RNP, *Arid1a*-RNP, *Pik3r1*-RNP, *Muc16*-RNP, *Kmt2d*-RNP and *Chd4*-RNP, oligonucleotides containing target sequences in tandem were used (CRISPR tandem assays). Oligonucleotides were obtained lyophilized at IDT and were resuspended at a final concentration of 10µM in nuclease-free water. To obtain double-chain DNA fragments, oligonucleotides were amplified through PCR reaction. Oligonucleotides sequences, primers and PCR conditions are detailed in Supplementary Table SM3 and Supplementary Table SM4. PCR amplicons were resolved in agarose gels and purified using the NucleoSpin Gel and PCR Clean-up kit (740609, Macherey-Nagel) according to manufacturer’s instructions and quantified with NanoPhotometer® N60 (IMPLEN). Amplicons were submitted to cleavage assay with all ribonucleoproteins in a 5µl reactions volume containing 0.5µl of 15µM Cas9-RNP and 1µl of 50ng/µl PCR product in Optimized Minimal Essential Medium (Opti-MEM) (31985070, Gibco). Cleavage assay was performed at 37°C for 1 hour and the reaction was stopped at 85°C for 5 minutes. In vitro CRISPR/Cas9 assays were resolved in 3% agarose gels.

### Intrauterine electroporation of Cas9-RNP complexes

In vivo intrauterine electroporation was performed as previously described^15^ with some modifications. In vivo experiments were conducted using mTmG mice, aged between 3 and 6 months. Following the assembly of sgRNA-Cas9 complexes for electroporation, RNPs were pooled at the same proportion and surgical procedures were performed as described below. Female mice were anesthetized with 2% isoflurane via inhalation and received intraperitoneal analgesia (Buprex) at a dose of 8ml/kg. The animals were placed in ventral position, and a midline incision was made through the skin and the abdominal wall. Uterine horns were exposed, and 3µl of RNP pool, mixed with Fast Green FCF (68724, Sigma-Aldrich) for visualization, were injected into one uterine horn using a 700 Series Microliter Syringe, 30G (80408, Hamilton). After injection, using a BTX830 square electroporator (BTX), 4 pulses of 50mV for 50msec spaced by 950msec were applied to the injected uterine horn. This protocol was repeated, opposing the orientation of tweezers and performed along the entire length of the uterine horn. The tweezers used were Platinum Tweezertrode, 5mm diameter (BTX). The contralateral uterine horn was left untreated to serve as internal control. Once electroporated, uterine horns were returned to the abdominal cavity and the incision was closed using a wound stapler AutoClip System (12020-09, Fine Scientific Tools, FST). Mice were sacrificed by cervical dislocation at different time points post-electroporation, depending on the experiment. Uteri were dissected and processed as required for each analytical procedure.

### Isolation and Culture of Epithelial Endometrial Cells

Isolation of epithelial endometrial cells was performed as previously described^18^. In brief, mice were euthanized by cervical dislocation and uteri were dissected. Uterine horns were cut into 3mm pieces and washed in Hanks Balanced Salt Sodium (HBSS) (14175-046, Gibco). Uterine fragments were digested with 1% trypsin (15090-046, Gibco) in HBSS for 1 hour at 4°C and for 45 minutes at room temperature. Dulbecco’s modified Eagle’s medium (DMEM) (41965-018, Gibco) with 10% of inactivated Fetal Bovine Serum (FBSi) (A52567-01, Gibco) was added to stop trypsin reaction. Then, with the edge of a razor blade, epithelial sheets were squeezed out of the uterine fragments. Epithelial sheets were washed twice with Phosphate Buffered Saline (PBS) and resuspended in 1ml of basal medium (DMEM-F/12 (11039-021, Gibco) with 1mM sodium pyruvate (11360-070, Gibco), 1% penicillin/streptomycin (15140-122, Gibco) and 0.1% amphotericin B (I15290-018, Gibco)). Epithelial sheets were mechanically disrupted in clusters of cells by pipetting 50 times with a 1ml tip. Clusters were diluted in basal medium supplemented with 2% dextran-coated charcoal-stripped serum (DCC) (A33821-01, Gibco) and plated into 96-well plates (black with micro-clear bottom) (655077, Greiner Bio One). When required, phase-contrast or fluorescence images were taken with Eclipse Ts2R microscope (Nikon), and quantification of GFP-positive cells was performed using Fiji software (version 2.16).

### NGS-Amplicon Sequencing Analysis

Genomic DNA (gDNA) for NGS-amplicon sequencing analysis was obtained from fresh electroporated uteri, from frozen tumors and from paraffin embedded tissues. In all cases, gDNA extraction was carried out using the NucleoSpin Tissue (740952, Macherey-Nagel). Sample preparation was performed according to the type of sample: for fresh electroporated tissue, uteri were dissected and opened longitudinally to expose epithelial cells; frozen tissues were cut into 5mm pieces; and 4 slices of 3µm from paraffin-embedded tissues were deparaffined in xylol. Then, samples were digested with proteinase K for 3 hours at 37°C and protocol for gDNA extraction was carried out following manufacturer’s instructions. Genomic DNA was amplified by PCRs flanking RNP-targeted region. Primer sequences and PCR conditions are specified in supplementary material (Supplementary Table SM5 and Supplementary Table SM6). PCR amplicons were resolved in agarose gels, purified using the NucleoSpin Gel and PCR Clean-up kit, following manufacturer’s indications, and quantified with NanoPhotometer® N60. Next Generation Sequencing (NGS)-Amplicon sequencing was performed by Genewiz (Azenta Life Sciences) company. Raw Fastq sequences were submitted to bioinformatic analysis to determine the presence of indels using Crispresso2^19,20^ and Cas-Analyzer^21^ online tools. Editing percentages for targeted genes across different samples were analyzed and normalized using GraphPad Prism (version 8.0.1). For each sample, the gene with the highest editing percentage was set to 100% and the gene with the lowest to 0%, with all other gene values linearly scaled between these two extremes. This normalization allowed for direct comparison of editing profiles between samples, independently of absolute editing frequency.

### In situ RNA hybridization and rolling circle amplification for detection of CRISPR/Cas9-induced mutations

For padlock probes design, a detailed graphic of the design is provided in Supplementary Figure S2A. In brief, each crRNA target was used for determining the mRNA-specific region, each sequence was split in 2 and integrated into the padlock backbone. All padlock probes sequences are detailed in Supplementary Table SM7 and were obtained from IDT as 5’-phosphorylated DNA oligos, a requirement for ligation. For sample preparation, multiwell plates containing fibroblasts or epithelial cells were fixed for 10 minutes with 4% paraformaldehyde (PFA) and, after three PBS washes, incubated in methanol at -20°C for at least 4 hours. Then, cells were post-fixed with 4% PFA and preconditioned with hybridization buffer, containing 6X Saline Sodium Citrate (SSC) and 10% formamide (205820010, Thermo scientific). Hybridization of the padlock probes was done at a final concentration of 200nM for each probe (if more than one), in hybridization buffer, for 10 minutes at 55°C and overnight at 45°C. Then, RCA primers (Supplementary Table SM8) were hybridized to padlock probes at a concentration of 200nM for 1 hour at 45°C. Samples were washed three times with 20% formamide in 2X SSC and twice with PBS and 0.1% Tween20 (PBS-T). For padlock probes ligation, samples were incubated for 2 hours at 37°C with ligation mix, containing 50% glycerol, 1X T4 RNA ligase Buffer (B0216L, New England Biolabs, NEB), 10µM ATP (R0441, Thermo Fisher Scientific), 0.2µg/µl Recombinant Albumin (B9200, NEB), 0.1U/µl Ribolock RNase inhibitor (EO0382, Thermo Fisher Scientific) and 0.5U/µl SplintR ligase (M0375L, NEB). To wash the samples, they were rinsed twice with PBS-T. Rolling Circle Amplification (RCA) was performed for 2 hours at 42°C using 0.5U/µl EquiPhi29 DNA polymerase (A39391, Thermo Fisher Scientific) in a reaction containing 1X EquiPhi29 buffer, 1mM dithiothreitol (DTT) and 1mM dNTPs. Again, samples were washed three times with 20% formamide in 2X SSC and twice with PBS-T. The visualization of the RCA products was performed through hybridization of detection probes and/or fluorescent-labeled probes (Supplementary Table SM9). Specific detection probes and fluorescent-labelled probes were prehybridized at 95°C for 2 minutes and cooled to room temperature at a final concentration of 200nM/each. Then, samples were incubated with the mixture of hybridized probes and Hoechst, in hybridization buffer, for 2 hours at 45°C. Finally, non-hybridized probes were removed with three wash steps of 20% formamide in 2X SSC and two of PBS-T. Samples were mounted in aqueous mounting medium Fluoromount-G (00-4958-02, Thermo Fisher Scientific). Visualization and image acquisition was performed in EVOS FL inverted fluorescence microscope (Invitrogen), for generic probes detection, or with AX NSPARK confocal microscope (Nikon), for fluorochrome code detection.

### Image processing and RNA spots quantification

Multichannel *.nd2 files, obtained from AX NSPARK confocal microscope, were converted to *.tiff files, without compression or scaling. Cell segmentation and spot segmentation for each channel (Atto488, Cy3, Atto647 or AlexaFluor750) were performed in CellProfiler (version 4.2.6)^22^, using standard pipelines to identify nuclei, cell outlines and RNA spots. The output was exported as *.csv files, with individual spot positions and associated parent cells. Downstream analysis was performed in R (version 4.5.1)^23^ using tidyverse package^24^. Raw *.csv files containing spot data for each channel were imported and cell identifiers mapped. A threshold of 8 pixels was used as the maximum distance for defining spot colocalization between channels within the same cell. For each channel pair, the script calculates the number of colocalized spots per cell, using Euclidean distances between spot coordinates. Counting was performed for individual (non-colocalized) spots per channel, as well as for colocalized spots for each channel pair. Genes were assigned to single channels or colocalizations according to probe design. The final quantification table per cell includes individual and colocalization counts for all gene targets. For visualization of gene expression, spots counts were binarized (detection=1, if at least one spot detected, otherwise 0) and a binary heatmap was generated with the pheatmap package^25^. The complete R script, including data preprocessing, colocalization, and heatmap generation is available on request.

### Tissue processing and immunohistochemistry analysis on paraffin sections

For histological analysis, mice were euthanized, and uteri were dissected, formalin-fixed overnight at 4°C and paraffin-embedded. Paraffin sections of 3µm were dried for 1 hour at 80°C, dewaxed in xylene, gradually rehydrated in ethanol and washed in PBS. Antigen retrieval was performed in EnVision FLEX high pH Solution (K8004, DAKO) for 20 minutes at 95C. Then, samples were incubated with 3% H_2_O_2_, for endogenous peroxidase blocking, and washed three times with PBS. Primary antibodies were incubated for 30 minutes at room temperature, washed with PBS and incubated with Horseradish Peroxidase (HRP)-conjugated or biotin-conjugated secondary antibodies and streptavidin-HRP, depending on the primary antibody. Finally, staining was visualized through reaction with EnVision Detection kit (K4065, DAKO) using diaminobenzidine (DAB) substrate. Slides were counterstained with Harris hematoxylin. Primary and secondary antibodies used for immunohistochemistry, and their dilutions, are detailed in Supplementary Table SM10. When required, quantification of positive cells or positive nuclei in immunohistochemical stained section was performed using QuPath (version 0.5.1)^26^. For nuclear markers (p53), automated cell detection algorithms were employed to identify and classify nuclei based on stain intensitu. Intensity thresholds were established to segment nuclei into categories (negative, weak, moderate and strong) according to DAB chromogen signal, Histoscore (H-Score) quantification was subsequently calculated by combining the percentage of nuclei in each intensity category using the formula: H-score = (1 × % weakly positive) + (2 × % moderately positive) + (3 × % strongly positive), resulting in a value ranging from 0 to 300. For cellular markers (GFP or p16), positive cell detection was optimized for nuclear or cellular localization and exported as the percentage of positive cells. For both p53 and p16, a normal endometrial tissue sample was analyzed as a reference. All analyses were visually validated, and thresholding parameters were kept consistent across samples to allow quantitative comparison.

### Shallow WGS

Genomic DNA (gDNA) from frozen tissues was extracted as previously described (see *NGS-Amplicon Sequencing Analysis*). Shallow Whole Genome Sequencing (WGS) was performed by Genewiz (Azenta Life Sciences) company. Raw FASTQ files were quality-checked using the fastqcr package in R^23,27^. Reads were aligned to GRCm38/mm10 using BWA-MEM (v0.7.17)^28^. PCR duplicates were marked with MarkDuplicates tool from GATK (v4.3.0.0)^29^. Genome-wide coverage profiles were generated with readCounter from the HMMcopy suite (v0.99.0) using a fixed bin size of 1 Mb^30^. Copy number alterations (CNAs) were estimated from shallow whole-genome sequencing (sWGS) using the ichorCNA algorithm^31^. CNA data generated by ichorCNA were imported into R and represented as GRanges and RaggedExperiment objects. Protein-coding gene annotations for Mus musculus were retrieved from Ensembl (release 100) using the AnnotationHub package^32^.

## Supporting information

Supplementary Results tables 2

Supplementary Result Tables 1

## DATA AVAILABILITY STATEMENT

Data Availability Statement: The datasets analyzed during the current study are available from the corresponding author upon reasonable request.

## ETHICS APPROVAL AND CONSENT TO PARTICIPATE

All procedures performed in this study followed the National Institute of Health Guide for the Care and Use of Laboratory Animals and were compliant with the guidelines of Universitat de Lleida.

## AUTHOR CONTRIBUTIONS

MV-S, DL-N, and XD designed the research and developed the project concept. MV-S, DL-N, and XD prepared the manuscript. JE, ME, DOP, ADR, XM-G, DL-N, and XD provided essential materials and resources necessary for conducting the research. MV-S, NB, RN, and XD performed experiments and collected the data. MV-S, NB, RN, DL-N, and XD analyzed the data and conducted statistical analyses. All authors contributed to data interpretation and critically revised the manuscript.

