## Supplementary figures and images for "Multiplexed CRISPR/Cas9 Editing of Tumor Suppressor Genes Recapitulates Molecular and Morphological Features of High-Risk Endometrial Cancer"

### Supplementary Figures

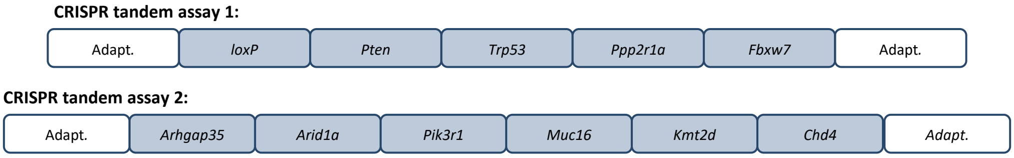

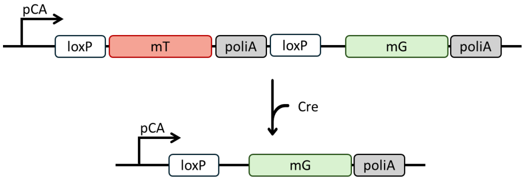

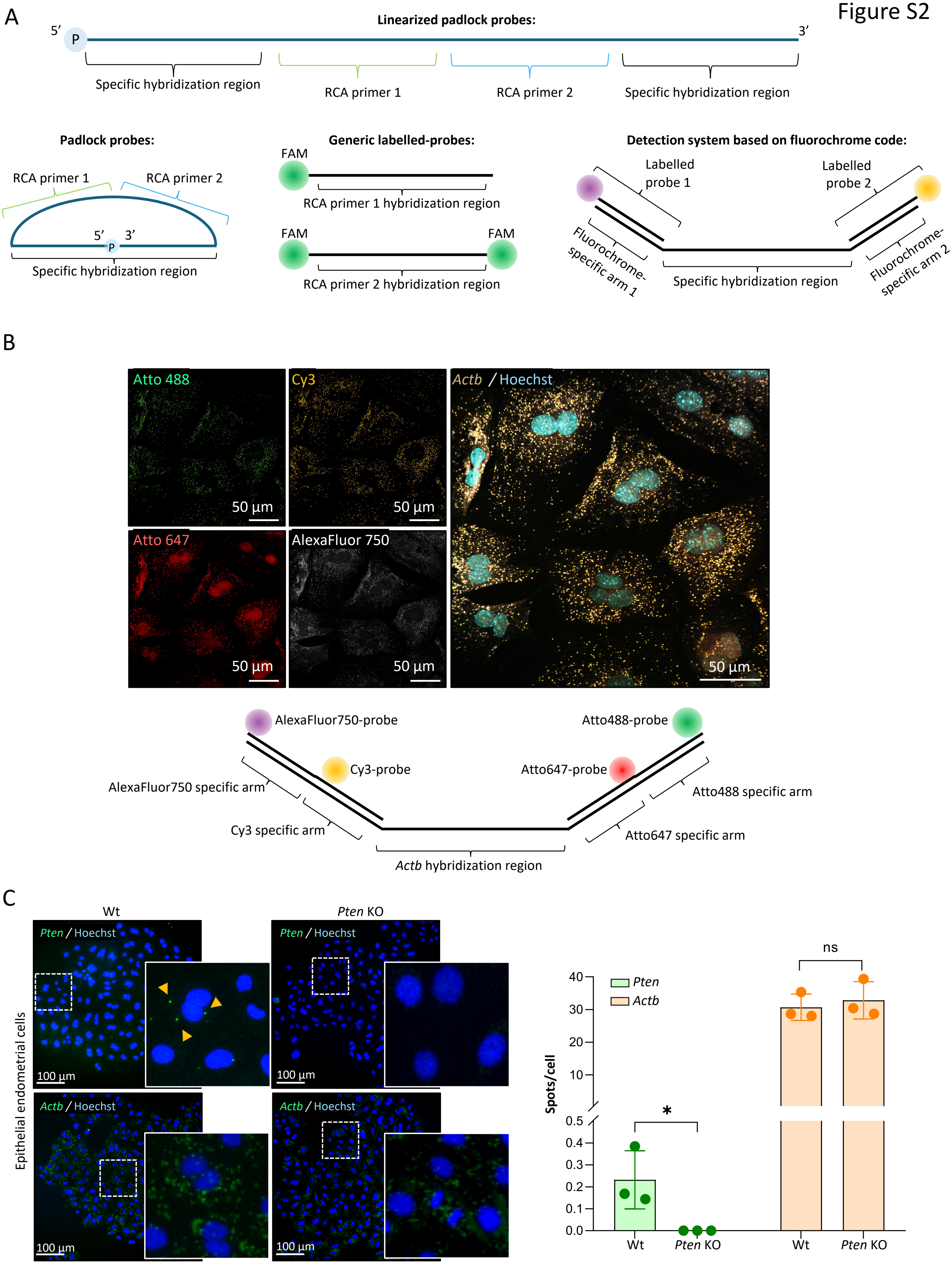

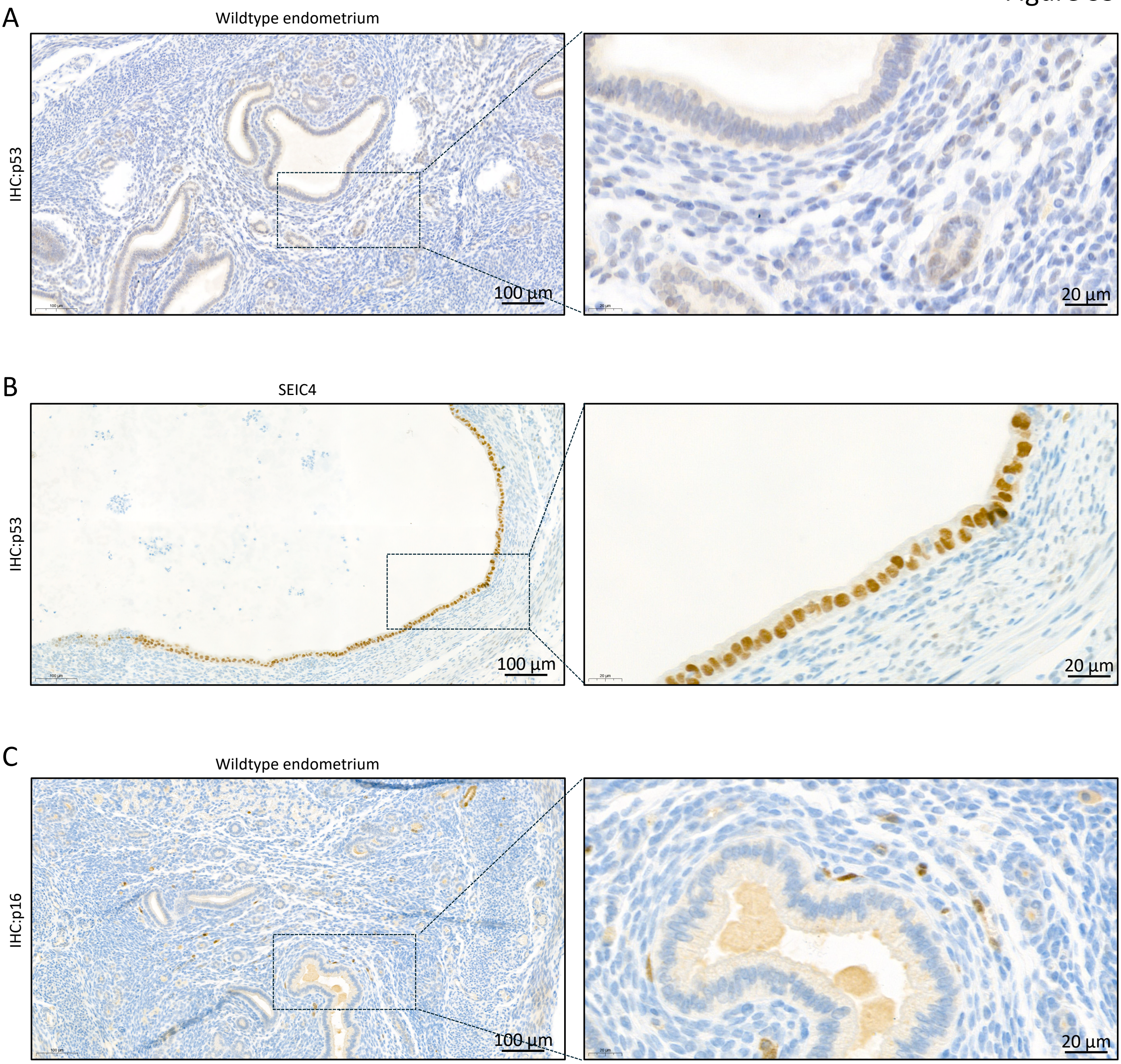
