## Supplementary Figure legends for "Multiplexed CRISPR/Cas9 Editing of Tumor Suppressor Genes Recapitulates Molecular and Morphological Features of High-Risk Endometrial Cancer"

**Figure S1. (A)** Representative diagram of DNA templates used for in vitro CRISPR/Cas9 tandem assay. **(B)** Structure of the pCA-mT/mG cassette, where pCA is Cytomegalovirus enhancer and chicken β-Actin promoter; polyA represent polyadenylation signals; mT encodes for the fluorescent protein tdTomato and mG the fluorescent protein EGFP.

**Figure S2. (A)** Illustration of the design of probes used for in situ RNA detection. Padlock probes, phosphorylated (P) at the 5' end, contain complementary regions of RCA primers 1 and 2 flanked by the specific hybridization region, complementary to the sgRNA target site. Generic detection probes are complementary to the RCA product at the padlock backbone and are labelled at 5' or at both 5' and 3' ends with FAM fluorochromes. Probes for the detection with fluorochrome code have a specific hybridization region to each gene and 1 or 2 arms complementary to the labelled probes. Labelled probes are marked with a molecule of Atto488, Cy3, Atto647 or AlexaFLuor750 at their 5' end. **(B)** *Actb* mRNA detection with the 4-colour probe, containing arms for all labelled probes, as represented in the lower panel. Images show detection with Atto488 (green), Cy3 (orange), Atto647 (red) and/or AlexaFluor750 (grey). Hoechst (blue) is used for nuclear staining. Scale bars: 50µm (60X magnification). **(C)** Left panel: representative images of *Pten* (green, upper) or *Actb* detection (green, lower) in epithelial endometrial cells from Cre:ER$^{T-/-}$ *Pten*$^{F/F}$ (Wt) and Cre:ER$^{T+/-}$ *Pten*$^{F/F}$ (*Pten* KO) mice. Orange arrows show *Pten* mRNA spots. Hoechst (blue) is used for nuclear staining. Scale bars: 100µm (100X magnification). Right panel: Spot quantification of samples showed in the left panel. Data is represented as mean of spots/cell ± s.e.m.

**Figure S3. (A)** Representative image of p53 immunohistochemistry (IHC) of sample SEIC4. Scale bars: 100µm (15X magnification) and 20µm (60X magnification). **(B)** Representative image of p53 immunohistochemistry (IHC) of wildtype endometrium, used as control for quantification of nuclear p53 and H-score in *Figure 5C*. Scale bars: 100µm (15X magnification) and 20µm (60X magnification). **(C)** Representative image of p16 immunohistochemistry (IHC) of wildtype endometrium, used as control for quantification of cytoplasmatic p16 in *Figure 5E*. Scale bars: 100µm (15X magnification) and 20µm (60X magnification).
