## Supplementary Results tables 2 for "Multiplexed CRISPR/Cas9 Editing of Tumor Suppressor Genes Recapitulates Molecular and Morphological Features of High-Risk Endometrial Cancer"

| ichorCNA | | |
|---|---|---|
| symbol | primari230609 | primari231129 |
| Hoxb1 | 1 | 3 |
| Hoxb2 | 1 | 3 |
| Hoxb3 | 1 | 3 |
| Skap1 | 1 | 3 |
| Hoxb4 | 1 | 3 |
| Caskin2 | 1 | 3 |
| Erbb2 | 1 | 3 |
| Grb7 | 1 | 3 |
| Hoxb5 | 1 | 3 |
| Ikzf3 | 1 | 3 |
| Llgl2 | 1 | 3 |
| Mien1 | 1 | 3 |
| Pgap3 | 1 | 3 |
| Tmem94 | 1 | 3 |
| Tsen54 | 1 | 3 |
| Cdk12 | 1 | 3 |
| Hoxb6 | 1 | 3 |
| Hoxb7 | 1 | 3 |
| Hoxb8 | 1 | 3 |
| Hoxb9 | 1 | 3 |
| Kbtbd8 | 3 | 1 |
| Neurod2 | 1 | 3 |
| Ormdl3 | 1 | 3 |
| Pnmt | 1 | 3 |
| Ppp1r1b | 1 | 3 |
| Stard3 | 1 | 3 |
| Suclg2 | 3 | 1 |
| Tcap | 1 | 3 |
| Zpbp2 | 1 | 3 |
| Acox1 | 1 | 3 |
| Arl6ip5 | 3 | 1 |
| Casc3 | 1 | 3 |
| Csf3 | 1 | 3 |
| Eif4e3 | 3 | 1 |
| Eogt | 3 | 1 |
| Fbf1 | 1 | 3 |
| Fbln2 | 3 | 1 |
| Fbxl20 | 1 | 3 |
| Foxp1 | 3 | 1 |
| Frmd4b | 3 | 1 |
| Gpr27 | 3 | 1 |
| Hoxb13 | 1 | 3 |
| Itga3 | 1 | 3 |
| Itgb4 | 1 | 3 |
| Lmod3 | 3 | 1 |
| Lrrc3c | 1 | 3 |
| Magi1 | 3 | 1 |
| Med1 | 1 | 3 |
| Mitf | 3 | 1 |
| Mrpl38 | 1 | 3 |
| Msl1 | 1 | 3 |
| Myo15b | 1 | 3 |
| Nr1d1 | 1 | 3 |

| | | |
|---|---|---|
| Pdk2 | 1 | 3 |
| Pdzrn3 | 3 | 1 |
| Prok2 | 3 | 1 |
| Rapgefl1 | 1 | 3 |
| Recql5 | 1 | 3 |
| Rybp | 3 | 1 |
| Sap30bp | 1 | 3 |
| Slc25a26 | 3 | 1 |
| Smim5 | 1 | 3 |
| Tafa1 | 3 | 1 |
| Ten1 | 1 | 3 |
| Thra | 1 | 3 |
| Tmf1 | 3 | 1 |
| Trim65 | 1 | 3 |
| Ttll6 | 1 | 3 |
| Uba3 | 3 | 1 |
| Wipf2 | 1 | 3 |
| Wnt7a | 3 | 1 |
| Abi3 | 1 | 3 |
| Adamts9 | 3 | 1 |
| Arl5c | 1 | 3 |
| B4galnt2 | 1 | 3 |
| Bptf | 1 | 3 |
| Cacnb1 | 1 | 3 |
| Calcoco2 | 1 | 3 |
| Cbx1 | 1 | 3 |
| Ccr7 | 1 | 3 |
| Cd300lg | 1 | 3 |
| Cdc42ep4 | 1 | 3 |
| Cdc6 | 1 | 3 |
| Cdk5rap3 | 1 | 3 |
| Cfap97d1 | 1 | 3 |
| Cntn4 | 3 | 1 |
| Copz2 | 1 | 3 |
| Cpsf4l | 1 | 3 |
| D11Wsu47e | 1 | 3 |
| Dhx8 | 1 | 3 |
| Dlx3 | 1 | 3 |
| Dlx4 | 1 | 3 |
| Etv4 | 1 | 3 |
| Exoc7 | 1 | 3 |
| Fam171a2 | 1 | 3 |
| Foxj1 | 1 | 3 |
| Fzd2 | 1 | 3 |
| Galk1 | 1 | 3 |
| Galr2 | 1 | 3 |
| Gjd3 | 1 | 3 |
| Gngt2 | 1 | 3 |
| Gpatch8 | 1 | 3 |
| Grb2 | 1 | 3 |
| Grn | 1 | 3 |
| Hdac11 | 3 | 1 |
| Igfbp4 | 1 | 3 |
| Itga2b | 1 | 3 |
| Kpnb1 | 1 | 3 |

| | | |
|---|---|---|
| Krt10 | 1 | 3 |
| Krt12 | 1 | 3 |
| Krt20 | 1 | 3 |
| Krt222 | 1 | 3 |
| Krt23 | 1 | 3 |
| Krt24 | 1 | 3 |
| Krt25 | 1 | 3 |
| Krt26 | 1 | 3 |
| Krt27 | 1 | 3 |
| Krt28 | 1 | 3 |
| Krt39 | 1 | 3 |
| Lrrc46 | 1 | 3 |
| Meox1 | 1 | 3 |
| Mpp3 | 1 | 3 |
| Mrpl10 | 1 | 3 |
| Nfe2l1 | 1 | 3 |
| Ngfr | 1 | 3 |
| Osbpl7 | 1 | 3 |
| Phb | 1 | 3 |
| Phospho1 | 1 | 3 |
| Plxdc1 | 1 | 3 |
| Pnpo | 1 | 3 |
| Prr15l | 1 | 3 |
| Psmd3 | 1 | 3 |
| Qrich2 | 1 | 3 |
| Rara | 1 | 3 |
| Rnf157 | 1 | 3 |
| Rpl19 | 1 | 3 |
| Rundc3a | 1 | 3 |
| Samd14 | 1 | 3 |
| Scrn2 | 1 | 3 |
| Shq1 | 3 | 1 |
| Slc25a39 | 1 | 3 |
| Slc4a1 | 1 | 3 |
| Smarce1 | 1 | 3 |
| Snx11 | 1 | 3 |
| Sost | 1 | 3 |
| Sp2 | 1 | 3 |
| Sp6 | 1 | 3 |
| Srp68 | 1 | 3 |
| Stac2 | 1 | 3 |
| Tafa4 | 3 | 1 |
| Tbkbp1 | 1 | 3 |
| Tbx21 | 1 | 3 |
| Tns4 | 1 | 3 |
| Top2a | 1 | 3 |
| Trim47 | 1 | 3 |
| Ubald2 | 1 | 3 |
| Unc13d | 1 | 3 |
| Unk | 1 | 3 |
| Wbp2 | 1 | 3 |
| Zfp652 | 1 | 3 |
| 1700001P01Rik | 1 | 3 |
| 1810010H24Rik | 1 | 3 |
| Aak1 | 3 | 1 |

| | | |
|---|---|---|
| Aanat | 1 | 3 |
| Abca5 | 1 | 3 |
| Abca6 | 1 | 3 |
| Abca8b | 1 | 3 |
| Abca9 | 1 | 3 |
| Abtb1 | 3 | 1 |
| Acbd4 | 1 | 3 |
| Acly | 1 | 3 |
| Adam11 | 1 | 3 |
| Add2 | 3 | 1 |
| Amz2 | 1 | 3 |
| Apoh | 1 | 3 |
| Arhgap23 | 1 | 3 |
| Arl4d | 1 | 3 |
| Armc7 | 1 | 3 |
| Arsg | 1 | 3 |
| Asb16 | 1 | 3 |
| Asprv1 | 3 | 1 |
| Atxn7l3 | 1 | 3 |
| Axin2 | 1 | 3 |
| Btbd17 | 1 | 3 |
| C87436 | 3 | 1 |
| Cacng1 | 1 | 3 |
| Cacng4 | 1 | 3 |
| Cacng5 | 1 | 3 |
| Ccdc43 | 1 | 3 |
| Cd300e | 1 | 3 |
| Cd300lb | 1 | 3 |
| Cdc27 | 1 | 3 |
| Cdr2l | 1 | 3 |
| Cep112 | 1 | 3 |
| Chchd4 | 3 | 1 |
| Chchd6 | 3 | 1 |
| Chl1 | 3 | 1 |
| Cntn3 | 3 | 1 |
| Cntn6 | 3 | 1 |
| Cog1 | 1 | 3 |
| Copg1 | 3 | 1 |
| Crbn | 3 | 1 |
| Crhr1 | 1 | 3 |
| Cwc25 | 1 | 3 |
| Cygb | 1 | 3 |
| Efcc1 | 3 | 1 |
| Eftud2 | 1 | 3 |
| Eif1 | 1 | 3 |
| Ern1 | 1 | 3 |
| Fads6 | 1 | 3 |
| Fam136a | 3 | 1 |
| Fbxo47 | 1 | 3 |
| Figla | 3 | 1 |
| Fkbp10 | 1 | 3 |
| G6pc3 | 1 | 3 |
| Gast | 1 | 3 |
| Gfpt1 | 3 | 1 |
| Gga3 | 1 | 3 |

| | | | |
|---|---|---|---|
| Gip | 1 | 3 | |
| Gjc1 | 1 | 3 | |
| Gm11639 | 1 | 3 | |
| Gmcl1 | 3 | 1 | |
| Gosr2 | 1 | 3 | |
| Gp9 | 3 | 1 | |
| Gpr142 | 1 | 3 | |
| Gpr179 | 1 | 3 | |
| Gprc5c | 1 | 3 | |
| Grin2c | 1 | 3 | |
| Gxylt2 | 3 | 1 | |
| H1f10 | 3 | 1 | |
| Hap1 | 1 | 3 | |
| Hdac5 | 1 | 3 | |
| Helz | 1 | 3 | |
| Hexim1 | 1 | 3 | |
| Hid1 | 1 | 3 | |
| Higd1b | 1 | 3 | |
| Hmces | 3 | 1 | |
| Hrob | 1 | 3 | |
| Igf2bp1 | 1 | 3 | |
| Il5ra | 3 | 1 | |
| Isy1 | 3 | 1 | |
| Itgb3 | 1 | 3 | |
| Itpr1 | 3 | 1 | |
| Jmjd6 | 1 | 3 | |
| Jpt1 | 1 | 3 | |
| Jup | 1 | 3 | |
| Kansl1 | 1 | 3 | |
| Kat7 | 1 | 3 | |
| Kcnj16 | 1 | 3 | |
| Kcnj2 | 1 | 3 | |
| Kctd2 | 1 | 3 | |
| Kif19a | 1 | 3 | |
| Klhl10 | 1 | 3 | |
| Klhl11 | 1 | 3 | |
| Krt14 | 1 | 3 | |
| Krt15 | 1 | 3 | |
| Krt16 | 1 | 3 | |
| Krt17 | 1 | 3 | |
| Krt19 | 1 | 3 | |
| Krt32 | 1 | 3 | |
| Krt34 | 1 | 3 | |
| Krt35 | 1 | 3 | |
| Krt36 | 1 | 3 | |
| Krt40 | 1 | 3 | |
| Krt9 | 1 | 3 | |
| Krtap16-1 | 1 | 3 | |
| Krtap3-1 | 1 | 3 | |
| Lasp1 | 1 | 3 | |
| Lrig1 | 3 | 1 | (HOMDEL en 1.8%) |
| Lsm12 | 1 | 3 | |
| Lsm3 | 3 | 1 | |
| Lyzl6 | 1 | 3 | |
| Map2k6 | 1 | 3 | |

| | | | |
|---|---|---|---|
| Mapt | 1 | 3 | |
| Mcm2 | 3 | 1 | |
| Med24 | 1 | 3 | (HOMDEL en 1.8%) |
| Meioc | 1 | 3 | |
| Mettl2 | 1 | 3 | |
| Mettl23 | 1 | 3 | |
| Mfsd11 | 1 | 3 | |
| Mgat5b | 1 | 3 | |
| Mgll | 3 | 1 | |
| Mif4gd | 1 | 3 | |
| Mpp2 | 1 | 3 | |
| Mrpl58 | 1 | 3 | |
| Mrps7 | 1 | 3 | |
| Mxd1 | 3 | 1 | |
| Myl4 | 1 | 3 | |
| Nags | 1 | 3 | |
| Nat9 | 1 | 3 | |
| Nbr1 | 1 | 3 | |
| Nfu1 | 3 | 1 | |
| Nmt1 | 1 | 3 | |
| Nol11 | 1 | 3 | |
| Npepps | 1 | 3 | |
| Nsf | 1 | 3 | |
| Nt5c | 1 | 3 | |
| Nt5c3b | 1 | 3 | |
| Nup210 | 3 | 1 | |
| Nup85 | 1 | 3 | |
| Nxph3 | 1 | 3 | |
| Otop2 | 1 | 3 | |
| Otop3 | 1 | 3 | |
| P3h4 | 1 | 3 | |
| Pcbp1 | 3 | 1 | |
| Pcyox1 | 3 | 1 | |
| Pecam1 | 1 | 3 | |
| Pip4k2b | 1 | 3 | |
| Pitpnc1 | 1 | 3 | |
| Plcd3 | 1 | 3 | |
| Plekhm1 | 1 | 3 | |
| Plxna1 | 3 | 1 | |
| Podxl2 | 3 | 1 | |
| Ppy | 1 | 3 | |
| Prcd | 1 | 3 | |
| Prickle2 | 3 | 1 | |
| Prkar1a | 1 | 3 | |
| Prkca | 1 | 3 | |
| Prpsap1 | 1 | 3 | |
| Psmd12 | 1 | 3 | |
| Pyy | 1 | 3 | |
| Rab37 | 1 | 3 | |
| Rab43 | 3 | 1 | |
| Rab7 | 3 | 1 | |
| Rad18 | 3 | 1 | |
| Rdm1 | 1 | 3 | |
| Rgs9 | 1 | 3 | |
| Rhbdf2 | 1 | 3 | |

| | | |
|---|---|---|
| Rpl23 | 1 | 3 |
| Rpl38 | 1 | 3 |
| Rpn1 | 3 | 1 |
| Rprml | 1 | 3 |
| Sdk2 | 1 | 3 |
| Slc16a5 | 1 | 3 |
| Slc16a6 | 1 | 3 |
| Slc25a19 | 1 | 3 |
| Slc39a11 | 1 | 3 |
| Smurf2 | 1 | 3 |
| Snf8 | 1 | 3 |
| Snrpg | 3 | 1 |
| Socs7 | 1 | 3 |
| Sox9 | 1 | 3 |
| Sphk1 | 1 | 3 |
| Sppl2c | 1 | 3 |
| Srgap3 | 3 | 1 |
| Srsf2 | 1 | 3 |
| Sstr2 | 1 | 3 |
| Ssu2 | 3 | 1 |
| St6galnac1 | 1 | 3 |
| St6galnac2 | 1 | 3 |
| Sumf1 | 3 | 1 |
| Sumo2 | 1 | 3 |
| Tac4 | 1 | 3 |
| Tex2 | 1 | 3 |
| Tgfa | 3 | 1 |
| Tia1 | 3 | 1 |
| Tmem101 | 1 | 3 |
| Tmem104 | 1 | 3 |
| Tmem106a | 1 | 3 |
| Tmem43 | 3 | 1 |
| Tmub2 | 1 | 3 |
| Tpra1 | 3 | 1 |
| Trnt1 | 3 | 1 |
| Ttyh2 | 1 | 3 |
| Ube2o | 1 | 3 |
| Ube2z | 1 | 3 |
| Ubtf | 1 | 3 |
| Ush1g | 1 | 3 |
| Wipi1 | 1 | 3 |
| Wnt3 | 1 | 3 |
| Wnt9b | 1 | 3 |
| Xpc | 3 | 1 |
| 1700012B07Rik | 1 | 3 |
| 1700123L14Rik | 3 | 1 |
| 1810020O05Rik | 3 | 1 |
| 2300003K06Rik | 1 | 3 |
| 4930590J08Rik | 3 | 1 |
| 4933428G20Rik | 1 | 3 |
| 9930022D16Rik | 1 | 3 |
| A730049H05Rik | 3 | 1 |
| Aarsd1 | 1 | 3 |
| Abca8a | 1 | 3 |
| Ace | 1 | 3 |

| | | |
|---|---|---|
| Ace3 | 1 | 3 |
| Aldh1l1 | 3 | 1 |
| Antxr1 | 3 | 1 |
| Anxa4 | 3 | 1 |
| Aoc2 | 1 | 3 |
| Aoc3 | 1 | 3 |
| Aplf | 3 | 1 |
| Arf2 | 1 | 3 |
| Arhgap25 | 3 | 1 |
| Arhgap27 | 1 | 3 |
| Arl8b | 3 | 1 |
| Atp5g1 | 1 | 3 |
| Atp5h | 1 | 3 |
| Atp6v0a1 | 1 | 3 |
| B230217C12Rik | 1 | 3 |
| BC048671 | 3 | 1 |
| Becn1 | 1 | 3 |
| Bhlhe40 | 3 | 1 |
| Bmp10 | 3 | 1 |
| Brca1 | 1 | 3 |
| C1ql1 | 1 | 3 |
| Cav3 | 3 | 1 |
| Cavin1 | 1 | 3 |
| Ccdc103 | 1 | 3 |
| Ccdc174 | 3 | 1 |
| Ccdc47 | 1 | 3 |
| Ccr10 | 1 | 3 |
| Cd300a | 1 | 3 |
| Cd300c | 1 | 3 |
| Cd300c2 | 1 | 3 |
| Cd300ld | 1 | 3 |
| Cd300ld2 | 1 | 3 |
| Cd300ld3 | 1 | 3 |
| Cd300ld4 | 1 | 3 |
| Cd300ld5 | 1 | 3 |
| Cd300lf | 1 | 3 |
| Cd79b | 1 | 3 |
| Cep95 | 1 | 3 |
| Cfap100 | 3 | 1 |
| Chst13 | 3 | 1 |
| Cisd3 | 1 | 3 |
| Cnbp | 3 | 1 |
| Cnp | 1 | 3 |
| Cntd1 | 1 | 3 |
| Cntnap1 | 1 | 3 |
| Coa3 | 1 | 3 |
| Coasy | 1 | 3 |
| Cyb561 | 1 | 3 |
| D6Ertd527e | 3 | 1 |
| Dcaf7 | 1 | 3 |
| Dcakd | 1 | 3 |
| Ddx42 | 1 | 3 |
| Ddx5 | 1 | 3 |
| Dhx58 | 1 | 3 |
| Dnaic2 | 1 | 3 |

| | | |
|---|---|---|
| Dnajb8 | 3 | 1 |
| Dnajc7 | 1 | 3 |
| Dusp3 | 1 | 3 |
| E030025P04Rik | 1 | 3 |
| Edem1 | 3 | 1 |
| Eefsec | 3 | 1 |
| Efcab15 | 1 | 3 |
| Efcab3 | 1 | 3 |
| Epop | 1 | 3 |
| Evpl | 1 | 3 |
| Ezh1 | 1 | 3 |
| Fam104a | 1 | 3 |
| Fam117a | 1 | 3 |
| Fam187a | 1 | 3 |
| Fam20a | 1 | 3 |
| Fdxr | 1 | 3 |
| Fgd5 | 3 | 1 |
| Fmnl1 | 1 | 3 |
| Ftsj3 | 1 | 3 |
| G6pc | 1 | 3 |
| Gata2 | 3 | 1 |
| Gfap | 1 | 3 |
| Gh | 1 | 3 |
| Ghdc | 1 | 3 |
| Gkn1 | 3 | 1 |
| Gkn2 | 3 | 1 |
| Gkn3 | 3 | 1 |
| Gm10234 | 3 | 1 |
| Gm11554 | 1 | 3 |
| Gm11555 | 1 | 3 |
| Gm11559 | 1 | 3 |
| Gm11562 | 1 | 3 |
| Gm11563 | 1 | 3 |
| Gm11564 | 1 | 3 |
| Gm11565 | 1 | 3 |
| Gm11567 | 1 | 3 |
| Gm11568 | 1 | 3 |
| Gm11569 | 1 | 3 |
| Gm11595 | 1 | 3 |
| Gm11596 | 1 | 3 |
| Gm11627 | 1 | 3 |
| Gm11634 | 1 | 3 |
| Gm11937 | 1 | 3 |
| Gm11938 | 1 | 3 |
| Gm11939 | 1 | 3 |
| Gm14180 | 1 | 3 |
| Gm14190 | 1 | 3 |
| Gm15737 | 3 | 1 |
| Gm20696 | 3 | 1 |
| Gm27029 | 1 | 3 |
| Gm44790 | 3 | 1 |
| Gm45140 | 3 | 1 |
| Gm765 | 3 | 1 |
| Gm884 | 1 | 3 |
| Gna13 | 1 | 3 |

| | | |
|---|---|---|
| Grip2 | 3 | 1 |
| Grm7 | 3 | 1 |
| Gsdma | 1 | 3 |
| Gsdma2 | 1 | 3 |
| Gsdma3 | 1 | 3 |
| H3f3b | 1 | 3 |
| Hcrt | 1 | 3 |
| Hexim2 | 1 | 3 |
| Hsd17b1 | 1 | 3 |
| Hspb9 | 1 | 3 |
| Icam2 | 1 | 3 |
| Ifi35 | 1 | 3 |
| Iqsec1 | 3 | 1 |
| Kat2a | 1 | 3 |
| Kbtbd12 | 3 | 1 |
| Kcnh4 | 1 | 3 |
| Kcnh6 | 1 | 3 |
| Kif18b | 1 | 3 |
| Klf15 | 3 | 1 |
| Kpna2 | 1 | 3 |
| Krt13 | 1 | 3 |
| Krt31 | 1 | 3 |
| Krt33a | 1 | 3 |
| Krt33b | 1 | 3 |
| Krt42 | 1 | 3 |
| Krtap1-3 | 1 | 3 |
| Krtap1-4 | 1 | 3 |
| Krtap1-5 | 1 | 3 |
| Krtap17-1 | 1 | 3 |
| Krtap2-4 | 1 | 3 |
| Krtap29-1 | 1 | 3 |
| Krtap3-2 | 1 | 3 |
| Krtap3-3 | 1 | 3 |
| Krtap31-1 | 1 | 3 |
| Krtap31-2 | 1 | 3 |
| Krtap4-1 | 1 | 3 |
| Krtap4-13 | 1 | 3 |
| Krtap4-16 | 1 | 3 |
| Krtap4-2 | 1 | 3 |
| Krtap4-6 | 1 | 3 |
| Krtap4-7 | 1 | 3 |
| Krtap4-8 | 1 | 3 |
| Krtap4-9 | 1 | 3 |
| Krtap9-1 | 1 | 3 |
| Krtap9-3 | 1 | 3 |
| Krtap9-5 | 1 | 3 |
| Limd2 | 1 | 3 |
| Lmcd1 | 3 | 1 |
| Lrrc37a | 1 | 3 |
| Lrrn1 | 3 | 1 |
| Map3k14 | 1 | 3 |
| Map3k3 | 1 | 3 |
| Marchf10 | 1 | 3 |
| Milr1 | 1 | 3 |
| Mllt6 | 1 | 3 |

| | | |
|---|---|---|
| Mlx | 1 | 3 |
| Mrc2 | 1 | 3 |
| Mrpl45 | 1 | 3 |
| Mrps25 | 3 | 1 |
| Mxra7 | 1 | 3 |
| Naglu | 1 | 3 |
| Nkiras2 | 1 | 3 |
| Nr2c2 | 3 | 1 |
| Oxtr | 3 | 1 |
| Pcgf2 | 1 | 3 |
| Plekhh3 | 1 | 3 |
| Polg2 | 1 | 3 |
| Ppp4r2 | 3 | 1 |
| Prokr1 | 3 | 1 |
| Prr29 | 1 | 3 |
| Psmb3 | 1 | 3 |
| Psmc3ip | 1 | 3 |
| Psmc5 | 1 | 3 |
| Psme3 | 1 | 3 |
| Ptges3l | 1 | 3 |
| Rab5c | 1 | 3 |
| Ramp2 | 1 | 3 |
| Rbsn | 3 | 1 |
| Retreg3 | 1 | 3 |
| Rnd2 | 1 | 3 |
| Rpl27 | 1 | 3 |
| Rundc1 | 1 | 3 |
| Ruvbl1 | 3 | 1 |
| Scn4a | 1 | 3 |
| Sec61a1 | 3 | 1 |
| Setmar | 3 | 1 |
| Slc35b1 | 1 | 3 |
| Slc41a3 | 3 | 1 |
| Slc6a6 | 3 | 1 |
| Slc9a3r1 | 1 | 3 |
| Smarcd2 | 1 | 3 |
| Smim6 | 1 | 3 |
| Snrnp27 | 3 | 1 |
| Spata32 | 1 | 3 |
| Spop | 1 | 3 |
| Srcin1 | 1 | 3 |
| Stat3 | 1 | 3 |
| Stat5a | 1 | 3 |
| Stat5b | 1 | 3 |
| Strada | 1 | 3 |
| Taco1 | 1 | 3 |
| Tanc2 | 1 | 3 |
| Tcam1 | 1 | 3 |
| Tlk2 | 1 | 3 |
| Trh | 3 | 1 |
| Trim80 | 1 | 3 |
| Ttc25 | 1 | 3 |
| Tubg1 | 1 | 3 |
| Tubg2 | 1 | 3 |
| Txnrd3 | 3 | 1 |

| | | |
|---|---|---|
| Uroc1 | 3 | 1 |
| V1ra8 | 3 | 1 |
| Vat1 | 1 | 3 |
| Vmn1r40 | 3 | 1 |
| Vmn1r41 | 3 | 1 |
| Vmn1r42 | 3 | 1 |
| Vmn1r43 | 3 | 1 |
| Vmn1r44 | 3 | 1 |
| Vmn1r45 | 3 | 1 |
| Vmn1r46 | 3 | 1 |
| Vmn1r47 | 3 | 1 |
| Vmn1r48 | 3 | 1 |
| Vmn1r49 | 3 | 1 |
| Vmn1r50 | 3 | 1 |
| Vmn1r51 | 3 | 1 |
| Vmn1r52 | 3 | 1 |
| Vmn1r53 | 3 | 1 |
| Vmn1r54 | 3 | 1 |
| Vps25 | 1 | 3 |
| Wnk4 | 1 | 3 |
| Zfp385c | 1 | 3 |
| Zxdc | 3 | 1 |
