## Supplementary Methods for "Multiplexed CRISPR/Cas9 Editing of Tumor Suppressor Genes Recapitulates Molecular and Morphological Features of High-Risk Endometrial Cancer"

**Supplementary Table SM1**. Genotyping primers and PCR conditions.

| Allele | | Primers | PCR conditions: Temp. | Time | Cycles | PCR product | |
|---|---|---|---|---|---|---|---|
| **Cre:ER$^{(T)}$** | Fwd: | ACGAACCTGGTCGAAATCGTGCG | 94 ºC | 2 min | 1 | | |
| | Rev: | CGGTCGATGCAACGAGTGATGAG | 94 ºC | 45 sec | 32 | Cre:ER$^{T-/-}$ | No band |
| | | | 65 ºC | 45 sec | | | |
| | | | 72 ºC | 45 sec | | | |
| | | | 72 ºC | 5 min | 1 | Cre:ER$^{T+/-}$ | 350 bp |
| | | | 4 ºC | ∞ | | | |
| ***Pten*$^{f/f}$** | Fwd: | CAAGCACTCTGCGAACTGAG | 94 ºC | 3 min | 1 | *Pten*$^{+/+}$ | 156 bp |
| | Rev: | AAGTTTTTGAAGGCAAGATGC | 94 ºC | 30 sec | 35 | | |
| | | | 60 ºC | 1 min | | *Pten*$^{f/+}$ | 156 bp & 328 bp |
| | | | 72 ºC | 2 min | | | |
| | | | 72 ºC | 2 min | 1 | *Pten*$^{f/f}$ | 328 bp |
| | | | 4 ºC | ∞ | | | |
| ***mTmG*** | Comú: | CTCTGCTGCCTCCTGGCTTCT | 94 ºC | 2 min | 1 | *mTmG*$^{+/+}$ | 330 bp |
| | Wt: | CGAGGCGGATCACAAGCAATA | 94 ºC | 30 sec | 35 | | |
| | Mut: | TCAATGGGCGGGGGTCGTT | 59 ºC | 30 sec | | *mTmG*$^{f/+}$ | 250 bp & 330 bp |
| | | | 72 ºC | 2 min | | | |
| | | | 72 ºC | 5 min | 1 | *mTmG*$^{f/f}$ | 250 bp |
| | | | 4 ºC | ∞ | | | |

**Supplementary Table SM2**. Sequence of crRNA targets used for CRISPR/Cas9 gene editing. Underlined nucleotides represent PAM sequence.

| Target gene | crRNA target |
|---|---|
| **mTmG *loxP*** | GTATGCTATACGAAGTTATTAGG |
| ***Fbxw7* (exon 5)** | GTTGTTGGTGTTGCTGAACATGG |
| ***Pten* (exon 5)** | AATTCACTGTAAAGCTGGAAAGG |
| ***Trp53* (exon 7)** | GGAGTCTTCCAGTGTGATGATGG |
| ***Ppp2r1a* (exon 5)** | TGTCATCTGAGCACAGGTTCCGG |
| ***Arhgap35* (exon 1)** | AAGACTTCCCAATGCCGCACTGG |
| ***Arid1a* (exon 1)** | AAGAACTCGAACGGGAACGCGGG |
| ***Pik3r1* (exon 1)** | CTTACGTTGAATACATTGGAAGG |
| ***Muc16* (exon 2)** | ATAAAGAGGGCTTCTCGTCAGGG |
| ***Kmt2d* (exon 4)** | ACTGCCAACTGGCACGCTTGCGG |
| ***Chd4* (exon 2)** | TCGGACCCTCACCAACTACAAGG |

**Supplementary Table SM3**. Sequence of DNA oligonucleotides used for CRISPR *in vitro* tandem assays. Amplification regions are marked in blue and gene targets are separated with slashes.

| | DNA sequence |
|---|---|
| **CRISPR *tandem assay* 1** | Amp fwd/*loxP*/*Fbxw7*/*Pten*/*Trp53*/*Ppp2r1a*/Amp rev |
| | GGAAGTGGCTCAGGTTCTGGA/GTATGCTATACGAAGTTATTAGG/AATTCACTGTAAAGCTGGAAAGG/GGAGTCTTCCAGTGTGATGATGG/TGTCATCTGAGCACAGGTTCCGG/CCATGTTCAGCAACACCAACAAC/CTCTGTATGCGATCGGCCAAG |
| **CRISPR *tandem assay* 2** | Amp fwd/*Arhgap35*/Arid1a/Pik3r1/Kmt2d/Chd4/Amp rev |
| | GGAAGTGGCTCAGGTTCTGGA/AAGACTTCCCAATGCCGCACTGG/AAGAACTCGAACGGGAACGCGGG/CTTACGTTGAATACATTGGAAGG/ATAAAGAGGGCTTCTCGTCAGGG/ACTGCCAACTGGCACGCTTGCGG/CCTTGTAGTTGGTGAGGGTCCGA/CTCTGTATGCGATCGGCCAAG |

**Supplementary Table SM4**. Sequence of primers and PCR conditions for CRISPR tandem assay oligonucleotides amplification.

| | Primer sequence | PCR program | | |
|---|---|---|---|---|
| | | **Temp.** | **Time** | **Cycles** |
| **CRISPR tandem assays 1, 2** | Fwd: AGAAGGAGATATAACTGGAAGTGGCTCAGGTTCTGGA<br>Rev: GGAGATGGGAAGTCACTTGGCCGATCGCATACAGAG | 96 ºC | 1 min | 1 |
| | | 96 ºC | 10 sec | |
| | | 65 ºC | 20 sec | 30 |
| | | 72 ºC | 30 sec | |
| | | 72 ºC | 10 min | 1 |
| | | 4 ºC | ∞ | |

**Supplementary Table SM5**. Sequence of primers used for NGS amplicon sequencing.

| **Target gene** (product size) | **Forward primer** | **Reverse primer** |
|---|---|---|
| ***Fbxw7*** (218 bp) | ATACATCTGGGGCAAGCTTAAGGTCTTACGTATAAGCAGG | CTGAAACATTTTCAGCCACTCTTGAAGGCCTGTGGGTGGT |
| ***Pten*** (230 bp) | ATCCTTTTGAAGACCATAACCCACCACAGCTAGAACTTAT | CTTTTTGTCTCTGGTCCTTACTTCCCCATAAAAATCTAGG |
| ***Trp53*** (220 bp) | GCCGGCTCTGAGTATACCACCATCCACTACAAGTACATGT | TAGGAACCAAAGAGCGTTGGGCATGTGGTAGGGGG |
| ***Ppp2r1a*** (225 bp) | AGTGCTCTGTGCTGTGCCGGGACACTAGCAGTTAGGACTT | CTGCTCGTCAGAGGCCAGGTTAGAGAACATGGGAATGATC |
| ***Arhgap35*** (245 bp) | ATGATGATGGCAAGAAAGCAAGATGTCCGCATCCCCACCT | GAGCGGCTAACTTCTCCCCAGTACAGAAAGTGGTCATTAT |
| ***Arid1a*** (479 bp) | CTCGGAGCTGAAGAAAGCCGAGCAGCAGCAGCGGGAGGA | GGCCAGGGCTTTGTTGTCCGCCATGTTGTTGGTGGAAGA |
| ***Pik3r1*** (200 bp) | CCAGGAAGCCCGGCCTGAAGATATTGGCTGGTTAAATGGC | CTTGCTGCTCCGTGTCAGCTTCAGTTTTTGAAGAACCCGG |
| ***Muc16*** (240 bp) | GAATTTTCACAGACAAGCTCTGCTTCCTCTGTTAACTCAG | ATTAGCTGTTCCAAAGATGGAGGTTAGCCCTATGTGTTCT |
| ***Kmt2d*** (235 bp) | AGGAGGCTCGCTGTGCAGTGTGTGAGGGGCCAGGGCAGCT | CAGGACACATTGTGGATCCTTCCTAGCCTCATTTTCTGTCCACAC |
| ***Chd4*** (256 bp) | TTAGAAGACTGGGGTATGGAAGACATCGACCATGTGTTCT | ATCTTGGAGACAGCAATCTTGGGGTTTTTAGCAGCAATCA |

**Supplementary Table SM6**. PCR conditions used for NGS amplicon sequencing.

| **PCR program** | | |
|---|---|---|
| **Temp.** | **Time** | **Cycles** |
| 95 ºC | 5 min | 1 |
| 95 ºC | 15 sec | 45 |
| 72 ºC | 30 sec | |
| 72 ºC | 5 min | 1 |
| 4 ºC | ∞ | 1 |

**Supplementary Table SM7.** Sequence of padlock probes used for RCA experiments. /5Phos/ indicates phosphorylation at 5' end.

| Target gene | Padlock probe sequence |
|---|---|
| ***Fbxw7*** | /5Phos/ACATGGTACAAGGCCAGTGGTTGACGTATATCTTGCGAGTGAGCGACCTCAATGCTGCTGCTGTACTACTAAAGTTGTTGGTGTTGCTGA |
| ***Pten*** | /5Phos/CAGCTTTACAGTGAATTGCTGCTGACGTATATCTTGCGAGTGAGCGACCTCAATGCTGCTGCTGTACTACTACACCAGTCCGTCCCTTTC |
| ***Trp53*** | /5Phos/TGATGGTAAGGATAGGTCGGCCTGACGTATATCTTGCGAGTGAGCGACCTCAATGCTGCTGCTGTACTACTCTGGAGTCTTCCAGTGTGA |
| ***Ppp2r1a*** | /5Phos/TTCCGGAAGTACTGTCGAAGCTGACGTATATCTTGCGAGTGAGCGACCTCAATGCTGCTGCTGTACTACTGGGTGTCATCTGAGCACAGG |
| ***Arhgap35*** | /5Phos/ACTGGCCTTTCTCCTTCTCACTGACGTATATCTTGCGAGTGAGCGACCTCAATGCTGCTGCTGTACTACTACAAGACTTCCCAATGCCGC |
| ***Arid1a*** | /5Phos/TTCCCGTTCGAGTTCTTCAGCTGACGTATATCTTGCGAGTGAGCGACCTCAATGCTGCTGCTGTACTACTGGGCGGGCCTAGGGCCCGCG |
| ***Pik3r1*** | /5Phos/AATGTATTCAACGTAAGTTCCTGACGTATATCTTGCGAGTGAGCGACCTCAATGCTGCTGCTGTACTACTGGTGAAATTCTTTTCCTTCC |
| ***Muc16*** | /5Phos/TCAGGGTTCTGCCAGCTCTACTGACGTATATCTTGCGAGTGAGCGACCTCAATGCTGCTGCTGTACTACTGGAATAAAGAGGGCTTCTCG |
| ***Kmt2d*** | /5Phos/TTGCGGGCAGTCAGAGCAGTCTGACGTATATCTTGCGAGTGAGCGACCTCAATGCTGCTGCTGTACTACTGGCACTGCCAACTGGCACGC |
| ***Chd4*** | /5Phos/AGTTGGTGAGGGTCCGATAACTGACGTATATCTTGCGAGTGAGCGACCTCAATGCTGCTGCTGTACTACTAAATTGGCTGAAGGCCTTGT |
| ***Actb*** | /5Phos/AGCACTTGCGGTGCACGATGCTGACGTATATCTTGCGAGTGAGCGACCTCAATGCTGCTGCTGTACTACTTCAGTAACAGTCCGCCTAGA |

**Supplementary Table SM8.** Sequence of RCA primers used in RCA experiments. Asterisks indicate phosphorothioate bonds.

| Primer | Primer sequence |
|---|---|
| **RCA 1** | ACAGCAGCAGCATTGAGG*T*C |
| **RCA 2** | CGCAAGATATACG*T*C |

**Supplementary Table SM9.** Sequence of detection probes used for visualization of RCA experiments.

| Target site (probe) | Probe sequence |
|---|---|
| **RCA 1** | /FAM/CCTCAATGCTGCTGCTGTACTAC |
| **RCA 2** | /FAM/CTGACGTATATCTTGCGAGTGAG/FAM/ |
| **Cy3-*T3*-Cy3** | /Cy3/GCGCGAAATTAACCCTCACTAAAGG/Cy3/ |
| **Atto488** | /ATTO488/TCCGAGTACGCT |
| **Cy3** | /Cy3/AGTCCGTAGCGA |
| **Atto647** | /ATTO647/CGTACGCTAGTC |
| **AlexaFluor750** | /AF750/TGCAGTCCGATC |
| ***Actb-T3*** | GTAACAGTCCGCCTAGAAGCACTTGCGGTGCACGACCTTTAGTGAGG GTTAATTTCGCGC |
| ***Actb*-4 color** | GATCGGACTGCATCGCTACGGACTGTAACAGTCCGCCTAGAAGCACT TGCGGTGGACTAGCGTACGAGCGTACTCGGA |
| ***Fbxw7*-Atto488** | TGTTGGTGTTGCTGAACATGGTACAAGGCCAGCGTACTCGGA |
| ***Pten*-Atto647** | AGTCCGTCCCTTTCCAGCTTTACAGTGAATGACTAGCGTACG |
| ***Trp53*-Cy3** | AGTCTTCCAGTGTGATGATGGTAAGGATAGTCGCTACGGACT |
| ***Ppp2r1a*-AF750** | TCATCTGAGCACAGGTTCCGGAAGTACTGTGATCGGACTGCA |
| ***Arhgap35*-Cy3/Atto647** | GACTAGCGTACGACTTCCCAATGCCGCACTGGCCTTTCTCCTTCGCT ACGGACT |
| ***Arid1a*-Atto488/Cy3** | AGCGTACTCGGAGGCCTAGGGCCCGCGTTCCCGTTCGAGTTCTCGCT ACGGACT |
| ***Pik3r1*-Atto488/AF750** | AGCGTACTCGGAAATTCTTTTCCTTCCAATGTATTCAACGTAGATCG GACTGCA |
| ***Muc16*-Cy3/AF750** | TCGCTACGGACTAAAGAGGGCTTCTCGTCAGGGTTCTGCCAGGATCG GACTGCA |
| ***Kmt2d*-Atto488/Atto647** | GACTAGCGTACGTGCCAACTGGCACGCTTGCGGGCAGTCAGAAGCGT ACTCGGA |
| ***Chd4*-Atto647/AF750** | GACTAGCGTACGGGCTGAAGGCCTTGTAGTTGGTGAGGGTCCGATCG GACTGCA |

**Supplementary Table SM10.** Antibodies used in immunohistochemistry analysis.

| Target protein | Dilution | Catalogue number, Company | Antigen Retrieval | Secondary Antibody |
|---|---|---|---|---|
| β-catenin | RTU | IR702, Dako | High pH | EV Flex Kit |
| **Cytokeratin 8** | RTU | Homemade Antibody | High pH | Rabbit-biotin |
| **GFP** | 1:100 | 600-101-215, Rockland | High pH | Goat-biotin |
| **p16** | RTU | CNIO Antibodies Core Unit | Unknown | Unknown |
| **p53** | RTU | CNIO Antibodies Core Unit | Unknown | Unknown |
| **Vimentin** | 1:500 | ab92547, Abcam | High pH | Rabbit-biotin |
| **Goat IgG-biotin** | 1:200 | SC-2489, Santacruz | | |
| **Rabbit IgG-biotin** | 1:200 | 111-065-144, Jackson | | |
| **Streptavidin-HRP** | 1:400 | P0397, Dako | | |
| **EnVision Flex Detection KIT** | RTU | K8002, Dako | | |
