## Supplementary Result Tables 1 for "Multiplexed CRISPR/Cas9 Editing of Tumor Suppressor Genes Recapitulates Molecular and Morphological Features of High-Risk Endometrial Cancer"

**Supplementary Table SR1.** Most frequent mutated TSG in high-risk endometrial cancer (cBioPortal).

| Gene | Samples | Mutation frequency | OncoKB recognition | OncoKB ONC/TSG | Described *in vivo* function in EC |
|---|---|---|---|---|---|
| ***TP53*** | 801 | 86.8% | Yes | TSG | Yes |
| ***PPP2R1A*** | 801 | 24.7% | Yes | TSG | Unknown |
| ***FBXW7*** | 801 | 19.4% | Yes | TSG | Yes |
| ***PTEN*** | 801 | 11.1% | Yes | TSG | Yes |
| ***ARID1A*** | 801 | 10.4% | Yes | TSG | Yes |
| ***PIK3R1*** | 801 | 12.1% | Yes | TSG | Unknown |
| ***KMT2D*** | 801 | 5% | Yes | TSG | Unknown |
| ***CHD4*** | 139 | 14.4% | Yes | Unknown | Unknown |
| ***ARHGAP35*** | 139 | 12.9% | Yes | TSG/ONC | Unknown |
| ***MUC16*** | 123 | 9.8% | No | Unknown | Unknown |

**Supplementary Table SR2.** Most frequent edits found 15 days post-electroporation of 10 TSG+LoxP.

| Fbxw7 **(exon 5)** | **Type** |
|---|---|
| TGGCCTTGTA**CCA**TGTTCAGCAACACCAAC | Ref |
| TGGCCTTGTACCATAATTCAGCAACACCAAC | Ins + Sub |
| Pten **(exon 5)** | **Type** |
| GCAGCAATTCACTGTAAAGCTGGAA**AGG**GA | Ref |
| GCAGCAATTCACTGTAAA----GAAAGGGA | Del |
| Trp53 **(exon 7)** | **Type** |
| CCTATCCTTA**CCA**TCATCACACTGGAAGAC | Ref |
| CCTATCCTTACCATCA----ACTGGAAGAC | Del |
| Ppp2r1a **(exon 5)** | **Type** |
| CAATCCCAGGTACTT**CCG**GAACCTGTGCTC | Ref |
| CAATCCCAGGTACTTCCGG-ACCTGTGCTC | Del |
| Arhgap35 **(exon 1)** | **Type** |
| AAAGG**CCA**GTGCGGCATTGGGAAGTCTTGT | Ref |
| AAAGGCCAGTG------TGGGAAGTCTTGT | Del |
| Arid1a **(exon 6)** | **Type** |
| CCTGAAGAACTCGAACGGGAACGC**GGG**CCCTAGGCCCGC | Ref |
| CCTGCAGAACTCGAACGGGAACGCGGGCCC | Sub |
| Pik3r1 **(exon 1)** | **Type** |
| TTGAATACATTGGA**AGG**AAAAGAATTTCAC | Ref |
| TTGAATACATTTGGAAGGAAAAGAATTTCAC | Ins |
| Muc16 **(exon 2** | **Type** |
| TGGCAGAA**CCC**TGACGAGAAGCCCTCTTTA | Ref |
| TGGCAGAACCCT---GAGAAGCCCTCTTTA | Del |
| Kmt2d **(exon 4)** | **Type** |
| TGACTGC**CCG**CAAGCGTGCCAGTTGGCAGT | Ref |
| TGACT----------------GTTGGCAGT | Del |
| Chd4 **(exon 2)** | **Type** |
| ACCCTCACCAACTACA**AGG**CCTTCAGCCAA | Ref |
| ACCCTCACC----ACAAGGCCTTCAGCCAA | Del |

**Supplementary Table SR3.** Summary of electroporated mice.

| **Sample ID** (intern code) | **Target genes** | **Birth date** | **Electroporation date** | **Disease progression** |
|---|---|---|---|---|
| **SEIC 1** (230529_5) | *LoxP, Pten, Trp53, Ppp2r1a, Fbxw7, Arhgap35, Arid1a, Pik3r1, Kmt2d, Chd4* | 28/9/2022 | 22/2/2023 | 13,71 weeks |
| **UCS 1** (230609_7) | *LoxP, Pten, Trp53, Ppp2r1a, Fbxw7, Arhgap35, Arid1a, Pik3r1, Kmt2d, Chd4* | 21/11/2022 | 9/3/2023 | 13,14 weeks |
| **UCS 2** (230704_5) | *LoxP, Pten, Trp53, Ppp2r1a, Fbxw7, Arhgap35, Arid1a, Pik3r1, Kmt2d, Chd4* | 21/11/2022 | 9/3/2023 | 16,71 weeks |
| **N 1** (230803_3) | *LoxP, Pten, Trp53, Ppp2r1a, Fbxw7, Arhgap35, Arid1a, Pik3r1, Kmt2d, Chd4* | 28/9/2022 | 22/2/2023 | 23,14 weeks |
| **N 2** (230803_4) | *LoxP, Pten, Trp53, Ppp2r1a, Fbxw7, Arhgap35, Arid1a, Pik3r1, Kmt2d, Chd4* | 28/9/2022 | 22/2/2023 | 23,14 weeks |
| **N 3** (230806_6) | *LoxP, Pten, Trp53, Ppp2r1a, Fbxw7* | 8/9/2022 | 30/11/2023 | 35,14 weeks |
| **UCS 3** (231002_6) | *LoxP, Pten, Trp53, Ppp2r1a, Fbxw7, Arhgap35, Arid1a, Pik3r1, Kmt2d, Chd4* | 21/11/2022 | 9/3/2023 | 29,57 weeks |
| **SEIC 2** (231002_14) | *LoxP, Pten, Trp53, Ppp2r1a, Fbxw7, Arhgap35, Arid1a, Pik3r1, Kmt2d, Chd4* | 20/12/2022 | 13/3/2023 | 29,00 weeks |
| **UCS 4** (231107_8) | *LoxP, Pten, Trp53, Ppp2r1a, Fbxw7, Arhgap35, Arid1a, Pik3r1, Kmt2d, Chd4* | 21/11/2022 | 9/3/2023 | 34,71 weeks |
| **SEIC 3** (231129_2) | *LoxP, Pten, Trp53, Ppp2r1a, Fbxw7* | 8/9/2022 | 30/11/2022 | 52,00 weeks |
| **UCS 5** (231129_4) | *LoxP, Pten, Trp53, Ppp2r1a, Fbxw7* | 8/9/2022 | 30/11/2022 | 52,00 weeks |
| **SEIC 4** (240201_9) | *LoxP, Pten, Trp53, Ppp2r1a, Fbxw7, Arhgap35, Arid1a, Pik3r1, Kmt2d, Chd4* | 20/12/2022 | 9/3/2023 | 47,00 weeks |
| **SEIC 5** (240201_13) | *LoxP, Pten, Trp53, Ppp2r1a, Fbxw7, Arhgap35, Arid1a, Pik3r1, Kmt2d, Chd4* | 20/12/2022 | 13/3/2023 | 46,43 weeks |
| **N 4** (240201_15) | *LoxP, Pten, Trp53, Ppp2r1a, Fbxw7, Arhgap35, Arid1a, Pik3r1, Kmt2d, Chd4* | 21/11/2022 | 9/3/2023 | 47,00 weeks |

**Supplementary Table SR4.** Most frequent edits in sample N1.

<table>
<tr><th colspan="4">N1</th></tr>
<tr><th>Fbxw7 (exon 5)</th><th>Type</th><th>% reads</th><th>% indels</th></tr>
<tr><td>TGGCCTTGTACCATGTTCAGCAACACCAAC</td><td>Ref</td><td>-</td><td>0</td></tr>
<tr><th>Pten (exon 5)</th><th>Type</th><th>% reads</th><th>% indels</th></tr>
<tr><td>GCAGCAATTCACTGTAAAGCTGGAAAGGGA</td><td>Ref</td><td>-</td><td rowspan="3">6.6</td></tr>
<tr><td>GCAGCAATTCACTGTAAAG-TGGAAAGGGA</td><td>Del</td><td>3.9</td></tr>
<tr><td>GCAGCAATTCACTGTAAAGCT-GAAAGGGA</td><td>Del</td><td>2.7</td></tr>
<tr><th>Trp53 (exon 7)</th><th>Type</th><th>% reads</th><th>% indels</th></tr>
<tr><td>CCTATCCTTACCATCATCACACTGGAAGAC</td><td>Ref</td><td>-</td><td rowspan="3">7.2</td></tr>
<tr><td>CCTATCCTTACCAT-----CACTGGAAGAC</td><td>Del</td><td>6.4</td></tr>
<tr><td>CCTATCCTTACCATCA-CACACTGGAAGAC</td><td>Del</td><td>0.8</td></tr>
<tr><th>Ppp2r1a (exon 5)</th><th>Type</th><th>% reads</th><th>% indels</th></tr>
<tr><td>CAATCCCAGGTACTTCCGGAACCTGTGCTC</td><td>Ref</td><td>-</td><td>0</td></tr>
<tr><th>Arhgap35 (exon 1)</th><th>Type</th><th>% reads</th><th>% indels</th></tr>
<tr><td>AAAGGCCAGTGCGGCATTGGGAAGTCTTGT</td><td>Ref</td><td>-</td><td rowspan="2">0.7</td></tr>
<tr><td>AAAGGCCAGT---GCATTGGGAAGTCTTGT</td><td>Del</td><td>0.7</td></tr>
<tr><th>Arid1a (exon 6)</th><th>Type</th><th>% reads</th><th>% indels</th></tr>
<tr><td>CTCGAACGGGAACGCGGGCCCTAGGCCCGC</td><td>Ref</td><td>-</td><td>0</td></tr>
<tr><th>Pik3r1 (exon 1)</th><th>Type</th><th>% reads</th><th>% indels</th></tr>
<tr><td>TTGAATACATTGGAAGGAAAAGAATTTCAC</td><td>Ref</td><td>-</td><td rowspan="3">4</td></tr>
<tr><td>TTGAATACATTATGGAAGGAAAAGAATTTCAC</td><td>Ins</td><td>1.7</td></tr>
<tr><td>TTGAATACA--GGAAGGAAAAGAATTTCAC</td><td>Del</td><td>1.5</td></tr>
<tr><th>Muc16 (exon 2</th><th>Type</th><th>% reads</th><th>% indels</th></tr>
<tr><td>TGGCAGAACCCTGACGAGAAGCCCTCTTTA</td><td>Ref</td><td>-</td><td>0</td></tr>
<tr><th>Kmt2d (exon 4)</th><th>Type</th><th>% reads</th><th>% indels</th></tr>
<tr><td>TGACTGCCCGCAAGCGTGCCAGTTGGCAGT</td><td>Ref</td><td>-</td><td rowspan="2">0.4</td></tr>
<tr><td>TGACTGCCCGCAATGCGTGCCAGTTGGCAGT</td><td>Ins</td><td>0.4</td></tr>
<tr><th>Chd4 (exon 2)</th><th>Type</th><th>% reads</th><th>% indels</th></tr>
<tr><td>ACCCTCACCAACTACAAGGCCTTCAGCCAA</td><td>Ref</td><td>-</td><td>0</td></tr>
</table>

**Supplementary Table SR5.** Most frequent edits in sample SEIC1.

| SEIC1 | | | |
|---|---|---|---|
| Fbxw7 **(exon 5)** | **Type** | **% reads** | **% indels** |
| TGGCCTTGTA**CCA**TGTTCAGCAACACCAAC | Ref | - | 0.6 |
| TGGCCTTGTACCATG-TCAGCAACACCAAC | Del | 0.2 | |
| Pten **(exon 5)** | **Type** | **% reads** | **% indels** |
| GCAGCAATTCACTGTAAAGCTGGAA**AGG**GA | Ref | - | 0 |
| Trp53 **(exon 7)** | **Type** | **% reads** | **% indels** |
| CCTATCCTTA**CCA**TCATCACACTGGAAGAC | Ref | - | 8.9 |
| CCTATCCTTACCATCA-CACACTGGAAGAC | Del | 1.8 | |
| CCTATCCTTACCATCA----ACTGGAAGAC | Del | 1.5 |  |
| Ppp2r1a **(exon 5)** | **Type** | **% reads** | **% indels** |
| CAATCCCAGGTACTT**CCG**GAACCTGTGCTC | Ref | - | 0.3 |
| CAATCCCAGGTACTTCCGGAA--------C | Del | 0.2 |  |
| CAATCCCAGGTACTTCCGGAA-CTGTGCTC | Del | 0.1 | |
| Arhgap35 **(exon 1)** | **Type** | **% reads** | **% indels** |
| AAAGG**CCA**GTGCGGCATTGGGAAGTCTTGT | Ref | - | 0 |
| Arid1a **(exon 6)** | **Type** | **% reads** | **% indels** |
| CTCGAACGGGAACGC**GGG**CCCTAGGCCCGC | Ref | - | 0.5 |
| CTCGAACGGG--CGCGGGCCCTAGGCCCGC | Del | 0.4 | |
| Pik3r1 **(exon 1)** | **Type** | **% reads** | **% indels** |
| TTGAATACATTGGA**AGG**AAAAGAATTTCAC | Ref | - | 3.8 |
| TTGAATACA-TGGAAGGAAAAGAATTTCAC | Del | 1.5 | |
| TTGAATAC---GGAAGGAAAAGAATTTCAC | Del | 0.1 | |
| Muc16 **(exon 2** | **Type** | **% reads** | **% indels** |
| TGGCAGAA**CCC**TGACGAGAAGCCCTCTTTA | Ref | - | 0 |
| Kmt2d **(exon 4)** | **Type** | **% reads** | **% indels** |
| TGACTGC**CCG**CAAGCGTGCCAGTTGGCAGT | Ref | - | 0 |
| Chd4 **(exon 2)** | **Type** | **% reads** | **% indels** |
| ACCCTCACCAACTACA**AGG**CCTTCAGCCAA | Ref | - | 0 |

**Supplementary Table SR6.** Most frequent edits in sample SEIC2.

| SEIC2 | | | |
|---|---|---|---|
| Fbxw7 **(exon 5)** | **Type** | **% reads** | **% indels** |
| TGGCCTTGTA**CCA**TGTTCAGCAACACCAAC | Ref | - | 1.2 |
| TGGCCTTGTACCATGTTCCAGCAACACCAAC | Ins | 0.6 | |
| TGGCCTTGTACCA---TCAGCAACACCAAC | Del | 0.4 | |
| Pten **(exon 5)** | **Type** | **% reads** | **% indels** |
| GCAGCAATTCACTGTAAAGCTGGAA**AGG**GA | Ref | - | 0.1 |
| GCAGCAATTCACTGTAAAGCTGG-AAGGGA | Del | 0.1 | |
| Trp53 **(exon 7)** | **Type** | **% reads** | **% indels** |
| CCTATCCTTA**CCA**TCATCACACTGGAAGAC | Ref | - | 3.2 |
| CCTATCCTTACCATCA-CACACTGGAAGAC | Del | 0.8 | |
| CCTATCCTTAC---CATCACACTGGAAGAC | Del | 0.4 | |
| Ppp2r1a **(exon 5)** | **Type** | **% reads** | **% indels** |
| CAATCCCAGGTACTT**CCG**GAACCTGTGCTC | Ref | - | 0 |
| Arhgap35 **(exon 1)** | **Type** | **% reads** | **% indels** |
| AAAGG**CCA**GTGCGGCATTGGGAAGTCTTGT | Ref | - | 0 |
| Arid1a **(exon 6)** | **Type** | **% reads** | **% indels** |
| CTCGAACGGGAACGC**GGG**CCCTAGGCCCGC | Ref | - | 0.4 |
| CTCGAACGGG-ACGCGGGCCCTAGGCCCGC | Del | 0.3 | |
| Pik3r1 **(exon 1)** | **Type** | **% reads** | **% indels** |
| TTGAATACATTGGA**AGG**AAAAGAATTTCAC | Ref | - | 1.2 |
| TTGAATACA-TGGAAGGAAAAGAATTTCAC | Del | 0.6 | |
| TTGAATACA--GGAAGGAAAAGAATTTCAC | Del | 0.3 | |
| Muc16 **(exon 2** | **Type** | **% reads** | **% indels** |
| TGGCAGAA**CCC**TGACGAGAAGCCCTCTTTA | Ref | - | 0 |
| Kmt2d **(exon 4)** | **Type** | **% reads** | **% indels** |
| TGACTGC**CCG**CAAGCGTGCCAGTTGGCAGT | Ref | - | 0 |
| Chd4 **(exon 2)** | **Type** | **% reads** | **% indels** |
| ACCCTCACCAACTACA**AGG**CCTTCAGCCAA | Ref | - | 0 |

**Supplementary Table SR7.** Most frequent edits in sample SEIC3.

| SEIC4 | | | |
|---|---|---|---|
| Fbxw7 **(exon 5)** | **Type** | **% reads** | **% indels** |
| TGGCCTTGTA**CCA**TGTTCAGCAACACCAAC | Ref | - | 4.5 |
| TGGCCTTGTACCATGTATCAGCAACACCAAC | Ins | 1.3 | |
| TGGCCTTGTACCATGTTATCAGCAACACCAAC | Ins | 0.3 | |
| Pten **(exon 5)** | **Type** | **% reads** | **% indels** |
| GCAGCAATTCACTGTAAAGCTGGAA**AGG**GA | Ref | - | 3.4 |
| GCAGCAATTCACTGTAA-----GAAAGGGA | Del | 2.1 |  |
| GCAGCAATTCACTGTAAAGC--GAAAGGGA | Del | 0.2 | |
| Trp53 **(exon 7)** | **Type** | **% reads** | **% indels** |
| CCTATCCTTA**CCA**TCATCACACTGGAAGAC | Ref | - | 6.7 |
| CCTATCCTTACCAT-----CACTGGAAGAC | Del | 3 |  |
| CCTATCCTTAC---CATCACACTGGAAGAC | Del | 1.2 | |
| Ppp2r1a **(exon 5)** | **Type** | **% reads** | **% indels** |
| CAATCCCAGGTACTT**CCG**GAACCTGTGCTC | Ref | - | 3 |
| CAATCCCAGGTACTTCCGG-ACCTGTGCTC | Del | 2.4 | |
| CAATCCCAGGTACTTCCGGAA--TGTGCTC | Del | 0.3 | |

**Supplementary Table SR8.** Most frequent edits in sample SEIC4.

| SEIC4 | | | |
|---|---|---|---|
| Fbxw7 **(exon 5)** | **Type** | **% reads** | **% indels** |
| TGGCCTTGTA**CCA**TGTTCAGCAACACCAAC | Ref | - | 0 |
| Pten **(exon 5)** | **Type** | **% reads** | **% indels** |
| GCAGCAATTCACTGTAAAGCTGGAA**AGG**GA | Ref | - | 0.2 |
| GCAGCAATTCACTGTAAAGCTGG-AAGGGA | Del | 0.1 | |
| GCAGCAATTCACTGTAA------GAAAGGGA | Del | 0.05 |  |
| Trp53 **(exon 7)** | **Type** | **% reads** | **% indels** |
| CCTATCCTTA**CCA**TCATCACACTGGAAGAC | Ref | - | 0 |
| Ppp2r1a **(exon 5)** | **Type** | **% reads** | **% indels** |
| CAATCCCAGGTACTT**CCG**GAACCTGTGCTC | Ref | - | 0 |
| Arhgap35 **(exon 1)** | **Type** | **% reads** | **% indels** |
| AAAGG**CCA**GTGCGGCATTGGGAAGTCTTGT | Ref | - | 0 |
| Arid1a **(exon 6)** | **Type** | **% reads** | **% indels** |
| CTCGAACGGGAACGC**GGG**CCCTAGGCCCGC | Ref | - | 0.4 |
| CTCGAACGGGAA---------CTAGGCCCGC | Del | 0.3 |  |
| Pik3r1 **(exon 1)** | **Type** | **% reads** | **% indels** |
| TTGAATACATTGGA**AGG**AAAAGAATTTCAC | Ref | - | 0.2 |
| TTGAATACA-TGGAAGGAAAAGAATTTCAC | Del | 0.1 | |
| TTGAATAC---GGAAGGAAAAGAATTTCAC | Del | 0.1 | |
| Muc16 **(exon 2** | **Type** | **% reads** | **% indels** |
| TGGCAGAA**CCC**TGACGAGAAGCCCTCTTTA | Ref | - | 0 |
| Kmt2d **(exon 4)** | **Type** | **% reads** | **% indels** |
| TGACTGC**CCG**CAAGCGTGCCAGTTGGCAGT | Ref | - | 0 |
| Chd4 **(exon 2)** | **Type** | **% reads** | **% indels** |
| ACCCTCACCAACTACA**AGG**CCTTCAGCCAA | Ref | - | 0.5 |
| ACCCTCACCAA-TACAAGGCCTTCAGCCAA | Del | 0.4 | |

**Supplementary Table SR9.** Most frequent edits in sample SEIC5.

| **SEIC5** | | | |
|---|---|---|---|
| Fbxw7 **(exon 5)** | **Type** | **% reads** | **% indels** |
| TGGCCTTGTA**CCA**TGTTCAGCAACACCAAC | Ref | - | |
| TGGCCTTGTACC----TCAGCAACACCAAC | Del | 3.1 | 11.3 |
| TGGCCTTGTACCATGT--------ACCAACAAC | Del | 2 |  |
| Pten **(exon 5)** | **Type** | **% reads** | **% indels** |
| GCAGCAATTCACTGTAAAGCTGGAA**AGG**GA | Ref | - | |
| GCAGCAATTCACTGTAAAGC--GAAAGGGA | Del | 1.6 | 3.8 |
| GCAGCAATTCACTGT-------GAAAGGGA | Del | 1.3 |  |
| Trp53 **(exon 7)** | **Type** | **% reads** | **% indels** |
| CCTATCCTTA**CCA**TCATCACACTGGAAGAC | Ref | - | |
| CCTATCCTTACCAT--TCACACTGGAAGAC | Del | 6 | 10.4 |
| CCTATCCTTAC---CATCACACTGGAAGAC | Del | 4.5 | |
| Ppp2r1a **(exon 5)** | **Type** | **% reads** | **% indels** |
| CAATCCCAGGTACTT**CCG**GAACCTGTGCTC | Ref | - | 0 |
| Arhgap35 **(exon 1)** | **Type** | **% reads** | **% indels** |
| AAAGG**CCA**GTGCGGCATTGGGAAGTCTTGT | Ref | - | |
| AAAGGCCAGTG-GGCATTGGGAAGTCTTGT | Del | 4.2 | 11 |
| AAAGGCCA--GCGGCATTGGGAAGTCTTGT | Del | 2.1 | |
| Arid1a **(exon 6)** | **Type** | **% reads** | **% indels** |
| CTCGAACGGGAACGC**GGG**CCCTAGGCCCGC | Ref | - | |
| CT----------CGCGGGCCCTAGGCCCGC | Del | 4.1 | 6.2 |
| CTCGAACGGGAAC-CGGGCCCTAGGCCCGC | Del | 2.1 | |
| Pik3r1 **(exon 1)** | **Type** | **% reads** | **% indels** |
| TTGAATACATTGGA**AGG**AAAAGAATTTCAC | Ref | - | |
| TTGAATACA-TGGAAGGAAAAGAATTTCAC | Del | 6.5 | 13.6 |
| TTGAATACATTTGGAAGGAAAAGAATTTCAC | Ins | 3 | |
| Muc16 **(exon 2** | **Type** | **% reads** | **% indels** |
| TGGCAGAA**CCC**TGACGAGAAGCCCTCTTTA | Ref | - | 0.3 |
| TGGCAGAACCC---CGAGAAGCCCTCTTTA | Del | 0.3 | |
| Kmt2d **(exon 4)** | **Type** | **% reads** | **% indels** |
| TGACTGC**CCG**CAAGCGTGCCAGTTGGCAGT | Ref | - | 0.2 |
| TGACT----------------GTTGGCAGT | Del | 0.2 |  |
| Chd4 **(exon 2)** | **Type** | **% reads** | **% indels** |
| ACCCTCACCAACTACA**AGG**CCTTCAGCCAA | Ref | - | |
| ACCCTCACCA---ACAAGGCCTTCAGCCAA | Del | 3 | 7.5 |
| ACCCTCACCA--TACAAGGCCTTCAGCCAA | Del | 2 | |

**Supplementary Table SR10.** Frequency of indel type for gene and tumor in samples SEIC1-5, expressed as % of reads.

| | | SEIC1 | SEIC2 | SEIC3 | SEIC4 | SEIC5 |
|---|---|---|---|---|---|---|
| *Fbxw7* | In-frame | 100% | 14.1% | 3.8% | 0% | 11% |
| | Frameshift | 0% | 85.9% | 96.2% | 0% | 89% |
| *Pten* | In-frame | 0% | 0% | 0% | 0% | 3% |
| | Frameshift | 0% | 100% | 100% | 100% | 97% |
| *Trp53* | In-frame | 12.5% | 16.7% | 22.3% | 0% | 33.4% |
| | Frameshift | 87.5% | 83.3% | 77.7% | 0% | 66.6 |
| *Ppp2r1a* | In-frame | 0% | 0% | 4.2% | 0% | 0% |
| | Frameshift | 100% | 0% | 95.8% | 100% | 0% |
| *Arhgap35* | In-frame | 0% | 0% | | 0% | 10% |
| | Frameshift | 0% | 0% | | 0% | 90% |
| *Arid1a* | In-frame | 0% | 0% | | 0% | 0% |
| | Frameshift | 100% | 100% | | 100% | 100% |
| *Pik3r1* | In-frame | 35.5% | 8.2% | | 40% | 6.5% |
| | Frameshift | 64.5% | 91.8% | | 60% | 93.5% |
| *Muc16* | In-frame | 0% | 0% | | 0% | 100% |
| | Frameshift | 0% | 0% | | 0% | 0% |
| *Kmt2d* | In-frame | 0% | 0% | | 0% | 0% |
| | Frameshift | 0% | 0% | | 0% | 100% |
| *Chd4* | In-frame | 0% | 0% | | 0% | 50% |
| | Frameshift | 0% | 0% | | 100% | 50% |

**Supplementary Table SR11** Most frequent edits in sample UCS1.

| **UCS1** | | | |
|---|---|---|---|
| Fbxw7 **(exon 5)** | **Type** | **% reads** | **% indels** |
| TGGCCTTGTA**CCA**TGTTCAGCAACACCAAC | Ref | - | 88.6 |
| TGGCCTTGTACCATAATTCAGCAACACCAAC | Ins + Sub | 30.5 | |
| TGGCCTTGTACCAT**G**TATCAGCAACACCAAC | Ins | 29.6 | |
| Pten **(exon 5)** | **Type** | **% reads** | **% indels** |
| GCAGCAATTCACTGTAAAGCTGGAA**AGG**GA | Ref | - | 80 |
| GCAGCAATTCACTGTAAA----GAAAGGGA | Del | 55.8 |  |
| Trp53 **(exon 7)** | **Type** | **% reads** | **% indels** |
| CCTATCCTTA**CCA**TCATCACACTGGAAGAC | Ref | - | 57.4 |
| CCTATCCTTACCATC-TCACACTGGAAGAC | Del | 19.2 | |
| CCTATCCTTAC---CATCACACTGGAAGAC | Del | 18.9 | |
| Ppp2r1a **(exon 5)** | **Type** | **% reads** | **% indels** |
| CAATCCCAGGTACTT**CCG**GAACCTGTGCTC | Ref | - | 0 |
| Arhgap35 **(exon 1)** | **Type** | **% reads** | **% indels** |
| AAAGG**CCA**GTGCGGCATTGGGAAGTCTTGT | Ref | - | 89.9 |
| AAAGGCCAGTG------TGGGAAGTCTTGT | Del | 32.1 |  |
| AAAGGCCAGTG-GGCATTGGGAAGTCTTGT | Del | 28.7 | |
| Arid1a **(exon 6)** | **Type** | **% reads** | **% indels** |
| CTCGAACGGGAACGC**GGG**CCCTAGGCCCGC | Ref | - | 93.8 |
| CTCGAACGGGAA--------CTAGGCCCGC | Del | 27.6 |  |
| CTCGAACGGGAAC-CGGGCCCTAGGCCCGC | Del | 25 | |
| Pik3r1 **(exon 1)** | **Type** | **% reads** | **% indels** |
| TTGAATACATTGGA**AGG**AAAAGAATTTCAC | Ref | - | 87 |
| TTGAATACA-TGGAAGGAAAAGAATTTCAC | Del | 58.4 | |
| Muc16 **(exon 2)** | **Type** | **% reads** | **% indels** |
| TGGCAGAA**CCC**TGACGAGAAGCCCTCTTTA | Ref | - | 37.2 |
| TGGCAGAACCCT---GAGAAGCCCTCTTTA | Del | 25.8 | |
| Kmt2d **(exon 4)** | **Type** | **% reads** | **% indels** |
| TGACTGC**CCG**CAAGCGTGCCAGTTGGCAGT | Ref | - | 52.6 |
| TGACT----------------GTTGGCAGT | Del | 33.3 |  |
| Chd4 **(exon 2)** | **Type** | **% reads** | **% indels** |
| ACCCTCACCAACTACA**AGG**CCTTCAGCCAA | Ref | - | 34.2 |
| ACCCTCACC----ACAAGGCCTTCAGCCAA | Del | 17 |  |
| ACCCTCACCA--TACAAGGCCTTCAGCCAA | Del | 7.1 | |

**Supplementary Table SR12.** Most frequent edits in sample UCS2.

| **UCS2** | | | |
|---|---|---|---|
| Fbxw7 **(exon 5)** | **Type** | **% reads** | **% indels** |
| TGGCCTTGTA**CCA**TGTTCAGCAACACCAAC | Ref | - | 70.5 |
| TGGCCTTGTACCATG-TCAGCAACACCAAC | Del | 43.3 | |
| TGGCCTTGTACCATGTT----AACACCAAC | Del | 2 |  |
| Pten **(exon 5)** | **Type** | **% reads** | **% indels** |
| GCAGCAATTCACTGTAAAGCTGGAA**AGG**GA | Ref | - | 84.7 |
| GCAGCAATTCACTGTAAAGCT-GAAAGGGA | Del | 51.9 | |
| Trp53 **(exon 7)** | **Type** | **% reads** | **% indels** |
| CCTATCCTTA**CCA**TCATCACACTGGAAGAC | Ref | - | 25.1 |
| CCTATCCTTACCATCA----ACTGGAAGAC | Del | 9.6 |  |
| CCTATCCTTACCATCA-CACACTGGAAGAC | Del | 5.9 | |
| Ppp2r1a **(exon 5)** | **Type** | **% reads** | **% indels** |
| CAATCCCAGGTACTT**CCG**GAACCTGTGCTC | Ref | - | 81.8 |
| CAATCCCAGGTACTTCCGGAA-CTGTGCTC | Del | 28.1 | |
| CAATCCCAGGTACTTCCGG-ACCTGTGCTC | Del | 25.9 | |
| Arhgap35 **(exon 1)** | **Type** | **% reads** | **% indels** |
| AAAGG**CCA**GTGCGGCATTGGGAAGTCTTGT | Ref | - | 91.3 |
| AAAGGCCA---CGGCATTGGGAAGTCTTGT | Del | 28.6 | |
| AAAGGCCA--GCGGCATTGGGAAGTCTTGT | Del | 23.8 | |
| Arid1a **(exon 6)** | **Type** | **% reads** | **% indels** |
| CTCGAACGGGAACGC**GGG**CCCTAGGCCCGC | Ref | - | 75.1 |
| CTCGAACGGGAATCGCGGGCCCTAGGCCCGC | Ins | 38.5 | |
| Pik3r1 **(exon 1)** | **Type** | **% reads** | **% indels** |
| TTGAATACATTGGA**AGG**AAAAGAATTTCAC | Ref | - | 69.5 |
| TTGAATACA-TGGAAGGAAAAGAATTTCAC | Del | 40.5 | |
| TTGAATAC-----------AAGAATTTCAC | Del | 1.5 |  |
| Muc16 **(exon 2** | **Type** | **% reads** | **% indels** |
| TGGCAGAA**CCC**TGACGAGAAGCCCTCTTTA | Ref | - | 0 |
| Kmt2d **(exon 4)** | **Type** | **% reads** | **% indels** |
| TGACTGC**CCG**CAAGCGTGCCAGTTGGCAGT | Ref | - | 0 |
| Chd4 **(exon 2)** | **Type** | **% reads** | **% indels** |
| ACCCTCACCAACTACA**AGG**CCTTCAGCCAA | Ref | - | 82.1 |
| ACCCTCACCAA--ACAAGGCCTTCAGCCAA | Del | 30.9 | |
| ACCCTCACCAA-TACAAGGCCTTCAGCCAA | Del | 25.9 | |

**Supplementary Table SR13** Most frequent edits in sample UCS3.

| UCS3 | | | |
|---|---|---|---|
| Fbxw7 **(exon 5)** | **Type** | **% reads** | **% indels** |
| TGGCCTTGTA**CCA**TGTTCAGCAACACCAAC | Ref | - | 1.9 |
| TGGCCTTGTACCATG-TCAGCAACACCAAC | Del | 0.8 | |
| TGGCCTTGTACCATGT-------CACCAAC | Del | 0.4 |  |
| Pten **(exon 5)** | **Type** | **% reads** | **% indels** |
| GCAGCAATTCACTGTAAAGCTGGAA**AGG**GA | Ref | - | 63.4 |
| GCAGCAATTCACTGTAAAGCT-GAAAGGGA | Del | 26.4 | |
| GCAGCAATTCACTGTAAAGC--GAAAGGGA | Del | 25.6 | |
| Trp53 **(exon 7)** | **Type** | **% reads** | **% indels** |
| CCTATCCTTA**CCA**TCATCACACTGGAAGAC | Ref | - | 32 |
| CCTATCCTTAC---CATCACACTGGAAGAC | Del | 32 | |
| Ppp2r1a **(exon 5)** | **Type** | **% reads** | **% indels** |
| CAATCCCAGGTACTT**CCG**GAACCTGTGCTC | Ref | - | 0 |
| Arhgap35 **(exon 1)** | **Type** | **% reads** | **% indels** |
| AAAGG**CCA**GTGCGGCATTGGGAAGTCTTGT | Ref | - | 79 |
| AAAGGCCAGTG-GGCATTGGGAAGTCTTGT | Del | 79 | |
| Arid1a **(exon 6)** | **Type** | **% reads** | **% indels** |
| CTCGAACGGGAACGC**GGG**CCCTAGGCCCGC | Ref | - | 89.5 |
| CTCGAACGGGAA--CGGGCCCTAGGCCCGC | Del | 78.9 | |
| Pik3r1 **(exon 1)** | **Type** | **% reads** | **% indels** |
| TTGAATACATTGGA**AGG**AAAAGAATTTCAC | Ref | - | 77.9 |
| TTGAATACA-TGGAAGGAAAAGAATTTCAC | Del | 60.5 | |
| Muc16 **(exon 2** | **Type** | **% reads** | **% indels** |
| TGGCAGAA**CCC**TGACGAGAAGCCCTCTTTA | Ref | - | 0 |
| Kmt2d **(exon 4)** | **Type** | **% reads** | **% indels** |
| TGACTGC**CCG**CAAGCGTGCCAGTTGGCAGT | Ref | - | 0 |
| Chd4 **(exon 2)** | **Type** | **% reads** | **% indels** |
| ACCCTCACCAACTACA**AGG**CCTTCAGCCAA | Ref | - | 80.3 |
| ACCCTCACCAAC-ACAAGGCCTTCAGCCAA | Del | 41.9 | |
| ACCCTCACCA--TACAAGGCCTTCAGCCAA | Del | 33.8 | |

**Supplementary Table SR14.** Most frequent edits in sample UCS4.

| UCS4 | | | |
|---|---|---|---|
| Fbxw7 **(exon 5)** | **Type** | **% reads** | **% indels** |
| TGGCCTTGTA**CCA**TGTTCAGCAACACCAAC | Ref | - | 12 |
| TGGCCTTGTACCATGTT-----ACACCAAC | Del | 5.1 |  |
| Pten **(exon 5)** | **Type** | **% reads** | **% indels** |
| GCAGCAATTCACTGTAAAGCTGGAA**AGG**GA | Ref | - | |
| GCAGCAATTCACTGTAA-----GAAAGGGA | Del | 5.7 | 15.5 |
| GCAGCAATTCACT----------AAAGGGA | Del | 2 |  |
| Trp53 **(exon 7)** | **Type** | **% reads** | **% indels** |
| CCTATCCTTA**CCA**TCATCACACTGGAAGAC | Ref | - | |
| CCTATCCTTACCAT-----CACTGGAAGAC | Del | 5.3 | 18.4 |
| CCTATCCTTAC---CATCACACTGGAAGAC | Del | 4 | |
| Ppp2r1a **(exon 5)** | **Type** | **% reads** | **% indels** |
| CAATCCCAGGTACTT**CCG**GAACCTGTGCTC | Ref | - | |
| CAATCCCAGGTACTTCCGGAA-CTGTGCTC | Del | 4 | 7.2 |
| CAATCCCAGGTACTTCCG---CCTGTGCTC | Del | 1.5 | |
| Arhgap35 **(exon 1)** | **Type** | **% reads** | **% indels** |
| AAAGG**CCA**GTGCGGCATTGGGAAGTCTTGT | Ref | - | |
| AAAGGCCAGT--GGCATTGGGAAGTCTTGT | Del | 6.5 | 11.8 |
| AAAG-------------------GTCTTGT | Del | 2.7 |  |
| Arid1a **(exon 6)** | **Type** | **% reads** | **% indels** |
| CTCGAACGGGAACGC**GGG**CCCTAGGCCCGC | Ref | - | |
| CTCGAACGGGAA---GGGCCCTAGGCCCGC | Del | 10.1 | 24.2 |
| CTCGAACGGGAA--CGGGCCCTAGGCCCGC | Del | 9.4 | |
| Pik3r1 **(exon 1)** | **Type** | **% reads** | **% indels** |
| TTGAATACATTGGA**AGG**AAAAGAATTTCAC | Ref | - | |
| TTGAATA----GGAAGGAAAAGAATTTCAC | Del | 10.4 | 15 |
| ---------TTGGAAGGAAAAGAATTTCAC | Del | 1.7 |  |
| Muc16 **(exon 2** | **Type** | **% reads** | **% indels** |
| TGGCAGAA**CCC**TGACGAGAAGCCCTCTTTA | Ref | - | 0.8 |
| TGGCAGAACC------AGAAGCCCTCTTTA | Del | 0.8 |  |
| Kmt2d **(exon 4)** | **Type** | **% reads** | **% indels** |
| TGACTGC**CCG**CAAGCGTGCCAGTTGGCAGT | Ref | - | 1.6 |
| TGACTGCCC----GCGTGCCAGTTGGCAGT | Del | 1.6 |  |
| Chd4 **(exon 2)** | **Type** | **% reads** | **% indels** |
| ACCCTCACCAACTACA**AGG**CCTTCAGCCAA | Ref | - | |
| ACCCTCACCAACT-CAAGGCCTTCAGCCAA | Del | 6.1 | 11.5 |
| ACCCTCACCA---ACAAGGCCTTCAGCCAA | Del | 1.4 | |

**Supplementary Table SR15.** Most frequent edits in sample UCS5.

<table>
<tr><th colspan="4">UCS5</th></tr>
<tr><th>Fbxw7 (exon 5)</th><th>Type</th><th>% reads</th><th>% indels</th></tr>
<tr><td>TGGCCTTGTACCATGTTCAGCAACACCAAC</td><td>Ref</td><td>-</td><td rowspan="2">53.8</td></tr>
<tr><td>TGGCCTTGTACCATG-TCAGCAACACCAAC</td><td>Del</td><td>20</td></tr>
<tr><th>Pten (exon 5)</th><th>Type</th><th>% reads</th><th>% indels</th></tr>
<tr><td>GCAGCAATTCACTGTAAAGCTGGAAAGGGA</td><td>Ref</td><td>-</td><td rowspan="3">45.8</td></tr>
<tr><td>GCAGCAATTCACTGTAAAGCT-GAAAGGGA</td><td>Del</td><td>30.3</td></tr>
<tr><td>GCAGCAATTCACTGTAAA----GAAAGGGA</td><td>Del</td><td>0.3</td></tr>
<tr><th>Trp53 (exon 7)</th><th>Type</th><th>% reads</th><th>% indels</th></tr>
<tr><td>CCTATCCTTACCATCATCACACTGGAAGAC</td><td>Ref</td><td>-</td><td rowspan="3">52.7</td></tr>
<tr><td>CCTATCCTTAC---CATCACACTGGAAGAC</td><td>Del</td><td>17.5</td></tr>
<tr><td>CCTATCCTTACCAT-----CACTGGAAGAC</td><td>Del</td><td>15.7</td></tr>
<tr><th>Ppp2r1a (exon 5)</th><th>Type</th><th>% reads</th><th>% indels</th></tr>
<tr><td>CAATCCCAGGTACTTCCGGAACCTGTGCTC</td><td>Ref</td><td>-</td><td rowspan="3">38.4</td></tr>
<tr><td>CAATCCCAGGTACTTCCGGAAGCCTGTGCTC</td><td>Ins</td><td>14</td></tr>
<tr><td>CAATCCCAGGTACTTCCGG-ACCTGTGCTC</td><td>Del</td><td>11.9</td></tr>
</table>

**Supplementary Table SR16.** Frequency of indel type for gene and tumor in samples UCS1-5, expressed as % of reads.

| | | UCS1 | UCS2 | UCS3 | UCS4 | UCS5 |
|---|---|---|---|---|---|---|
| *Fbxw7* | In-frame | 0% | 0% | 3.6% | 0% | 0% |
| | Frameshift | 100% | 100% | 96.4% | 100% | 100% |
| *Pten* | In-frame | 0% | 0% | 0% | 1% | 0% |
| | Frameshift | 100% | 100% | 100% | 99% | 100% |
| *Trp53* | In-frame | 47.6% | 7.4% | 100% | 27.3% | 55.2% |
| | Frameshift | 52.4% | 92.6% | 0% | 72.7% | 44.8% |
| *Ppp2r1a* | In-frame | 0% | 0% | 0% | 33.4% | 2.2% |
| | Frameshift | 0% | 100% | 0% | 66.6% | 97.8% |
| *Arhgap35* | In-frame | 56.7% | 49.4% | 0% | 0% | |
| | Frameshift | 45.3% | 50.6% | 100% | 100% | |
| *Arid1a* | In-frame | 8.6% | 70% | 0.6% | 48.4% | |
| | Frameshift | 91.4% | 30% | 99.4% | 51.6% | |
| *Pik3r1* | In-frame | 0% | 0.7% | 0% | 11.8% | |
| | Frameshift | 100% | 99.3% | 100% | 88.2% | |
| *Muc16* | In-frame | 100% | 0% | 0% | 100% | |
| | Frameshift | 0% | 0% | 0% | 0% | |
| *Kmt2d* | In-frame | 0% | 0% | 0% | 0% | |
| | Frameshift | 100% | 0% | 0% | 100% | |
| *Chd4* | In-frame | 0% | 2% | 0% | 13.3% | |
| | Frameshift | 100% | 98% | 100% | 86.7% | |
